# Microbial community composition, but not diversity, influence microbial necromass mineralization

**DOI:** 10.64898/2026.07.09.737581

**Authors:** Emmy L’Espérance, Vincent Poirier, Étienne Yergeau

## Abstract

Soil harbours a wide diversity of microbes responsible for essential functions, such as depolymerizing the C and N in organic matter through the production of exoenzymes. Some of these exoenzymes are universal, whereas others are specific to certain microbes. We hypothesized that higher microbial alpha diversity is associated with greater depolymerization capacity, specifically for protein and cellulose depolymerization, which will result in more N being mineralized. We therefore diluted two soil microbial communities, one from a forest soil and one from an agricultural soil, to create a diversity gradient. After nine weeks, we transferred these communities to a synthetic soil in which microbial necromass was the only nitrogen source. Before the transfer and two weeks after, we quantified protease, deaminase and β-glucosidase potential activity, characterized the bacterial and fungal communities, and measured the quantity of nitrogen mineralized. The dilution had very little effect on the processes measured, with no clear trend. For identical alpha diversity values, some communities had high process rates, while other not. It appeared that these communities varied widely, a side effect of the dilution approach, and that this variation was significantly linked to process rates. This shows that community composition (beta diversity) is more strongly related to enzymatic potential and mineralization than species richness (alpha diversity) following necromass addition. In conclusion, the relationship between diversity and depolymerization of microbial necromass is not simply a matter of a linear decrease along with diversity but is rather linked to how reduced diversity results in more stochastic microbial communities.

**Highlights:**

– Community composition (beta diversity) influence more microbial necromass depolymerization than species richness
– Abundance of specific microbes explained ammonification and nitrification processes
– Mineralization rates is different between crop and forest soil

## 1. Introduction

Many ecosystem processes, such as carbon (C) and nitrogen (N) cycling, are carried out by soil microbes. In some cases, such as nitrification, these processes are performed by well-defined narrow phylogenetic groups, whereas in other cases, such as N ammonification, a wide range of microorganisms are involved. In this latter case, since many soil organisms can perform the same function, there is functional redundancy (Heemsbergen et al., 2004; Chen et al., 2022), meaning that the loss of a microbial species does not always imply a loss of function (Loreau, 2004). Therefore, functional redundancy raises many questions regarding the role of microbial diversity in ecosystem functioning.

The link between microbial diversity and functions is better known for narrow processes such as nitrification and denitrification. On the one hand, nitrification is a two-step process where ammonium is oxidized into nitrate by restricted groups of microbes: ammonia-oxidizing bacteria and archaea, nitrite-oxidizing bacteria, and comammox (Shao et al., 2025). The presence of microbes belonging to these groups is crucial; without them, nitrification cannot occur. Accordingly, King et al. (2023) found that the loss of microbial alpha diversity results in a loss of nitrification (King et al., 2023). On the other hand, denitrification is the step-by-step reduction of nitrate into dinitrogen with different intermediates (i.e. NO_2_^-^, NO, N_2_O). Each step is performed by specific enzymes, and not all denitrifiers can perform complete denitrification (i.e. the reduction of nitrate to dinitrogen). Philippot et al. (2013) showed that even a small reduction in soil microbial alpha diversity significantly reduced denitrification rates by a factor of four to five (Philippot et al., 2013). Therefore, the presence of specific microbes is essential for complete denitrification, further proving that narrow processes are alpha diversity-dependent (Stein and Klotz, 2016; Pold et al., 2025). In contrast, broad processes are carried out by a diverse array of microbes, making them independent of the microbial diversity. However, among these broad processes, some of them are seen as a single process, but they grouped multiple distinct functions. Because they are perceived as a whole, these processes are often assumed to not be driven by microbial diversity, when in fact they could be. A good example is the depolymerization of the soil organic matter (SOM).

Nutrient release from soil organic matter in forms available for plant and microbial uptake involves diverse microbes. Among these bioavailable nutrients, there are C sources (sugars) and N sources (ammonium, nitrate and amino acids). The transformation of organic N into ammonium (N mineralization) is the result of two broad processes: depolymerization and ammonification. Mineralization is performed by a wide variety of heterotrophic microbes that use various molecules as energy (C) and nitrogen (N) sources. When their N demand is met, they will release excess N as ammonium (Wang et al., 2018). The limiting step of N mineralization, which is essential for replenishing soil N pools, is depolymerization (Schimel and Bennett, 2004). Taken as a whole, depolymerization is a grouped process, which can be defined as the breakdown of different complex organic compounds, like proteins, into simpler molecules, like amino acids, mediated by different enzymes (Jan et al., 2009). Proteins are the main source of organic N in the soil, coming from dead plants and microbial material (plant and microbial necromass), and making about 40% of the soil organic matter (Schulten and Schnitzer, 1997; Wanek et al., 2010).

Proteases are one of the main exoenzymes depolymerizing proteins, cleaving proteins into peptides and amino acids. Peptides and amino acids can be used by microbes and plants, or they can be further cleaved by microbial deaminases, releasing an amine. Due to a high diversity of N organic sources, N depolymerization is performed by a broad group of heterotrophic microbes, capable of producing a wide variety of proteases, deaminases and other exoenzymes (Sinsabaugh, 1994). Proteases can be classified based on their substrates and catalytic reactions, and some groups are associate to specific microbes, leading us to think that protease activity in the soil can be linked to microbial community alpha diversity and composition (Nguyen et al., 2019).

N depolymerization is closely associated with C depolymerization, since many of the N organic compounds contain a large amount of C. In the soil, microbial necromass (i.e., non-living microbes) represents around 50% of the total soil organic carbon (Wang et al., 2021). Microbial necromass also contains nitrogen in the form of chitin, protein, and peptidoglycan that needs to be depolymerized into amino sugars and amino acids (Buckeridge et al., 2022). Therefore, microbial necromass depolymerization encompasses multiple functions, mediated by different exoenzymes, linking N and C cycling. Some of them are catalyzed by exoenzymes universally produced, and others are more niche (Waldrop et al., 2000). Some studies have tried to define the relationship between plant litter depolymerization and microbial diversity. Baumann et al. (2013) manipulated microbial diversity and composition and found that reducing alpha diversity also reduced the breakdown of C^13^-labelled plant litter (Baumann et al., 2013). Plant litter contains mainly cellulose, depolymerized by β-glucosidase. Therefore, their study suggests that β-glucosidase activity is driven by microbial alpha diversity. On the other hand, microbial necromass is chemically distinct from plant necromass, and it is still unknown how microbial alpha diversity will affect microbial necromass mineralization and N depolymerization related to proteases and deaminases activities.

Microbial communities from cropland are usually more diverse than forest (Yoon et al., 2024). Microbial community composition and structure are also different between both ecosystems, because they are exposed to different nutrient inputs and to different ecosystems characteristic. Croplands are usually fertilized with inorganic N fertilizer, disturbing N cycling and microbial communities (Giller et al., 1997). The same way, different agriculture management practice, like tillage and pesticide application will alter microbial communities (Young and Ritz, 2000). Since forest are not disturbed by anthropological uses, microbial communities are used to nutrient coming from plant litter and have a higher fungal abundance than cropland (i.e., a higher fungal: bacterial ratio) (Szoboszlay et al., 2017). Since crop and forest soil microbial communities are distinct, depolymerization process could be differentially altered by microbial alpha diversity.

Since the link between microbial alpha diversity and broad grouped processes, such as depolymerization, is unclear, we quantified the depolymerization capacity of serially diluted crop and forest soils. Serial soil dilution was used to creates a diversity gradient of microbial communities, since dilution is associated with the loss of rare species among microbial communities (Dıaz et al., 2003; Philippot et al., 2013). We hypothesized that reducing microbial alpha diversity reduces C and N depolymerization (protease, deaminase and β-glucosidase), and N mineralization.

## 2. Materials and methods

### 2.1 Experimental design

We conducted a controlled experiment in Magenta boxes with two soil types (crop and forest soils) from Abitibi-Témiscamingue (Québec, Canada). Briefly, the crop soil contained: 0.023 g C g^-1^ soil and 0.00042 g N g^-1^ soil, C: N ratio= 54.76 and the forest soil contained: 0.022 g C g^-1^ soil and 0.0016 g N g^-1^ soil, C: N ratio= 13.37). The crop soil had a pH of 7.65 and the forest soil had a pH of 4.95 (measured in a 1:2 soil to CaCl_2_, 0.01 M ratio). This experiment was divided into two steps, where we had 6 levels of dilutions (D0, D2, D4, D6, D8 and a control), 2 soil types and 5 replicates, for a total of 60 experimental units. Firstly, we inoculated microbial dilutions into their native soil (crop and forest soil) and waited 9 weeks before quantifying enzymatic capacities, inorganic N content and sequencing microbial communities. Secondly, after 9 weeks, we used soil from each box to create microbial inocula and inoculated them into synthetic soil containing microbial necromass. The resulting boxes containing synthetic soil, microbial necromass and microbial inocula were incubated for two weeks before quantifying enzymatic capacities, inorganic N content and sequencing microbial communities.

### 2.2 Native soil

#### 2.2.1 Native soil preparation

We sieved both soil type at 4 mm and we reserved about 50 g of each soil to prepare microbial dilutions. The rest of the soil was air-dried, autoclaved and separated into 30 magenta boxes for each soil type (i.e., 50 g of dried soil per box).

#### 2.2.2 Microbial dilution preparation

We used the equivalent of 20 g of dry soil in 100 mL of Milli-Q water to create our D0 dilution solution (0.2 g soil mL^-1^ water). We blended D0 dilutions into a sterilized blender for 5 min at maximum speed and put them into a sterilized bottle. To create D1 dilution, we mixed 10 mL of the D0 solution into 90 mL of Milli-Q water. To create D2 dilution, we mixed 10 mL of the D1 solution into 90 mL of Milli-Q water and so on to make 9 dilutions. We inoculated 2.5 mL of D0, D2, D4, D6, and D8 into magenta boxes containing the autoclaved soil. We added water to keep every box at 70 % of the water-holding capacity of each soil type. After 9 weeks, we quantified potential enzymatic activities and inorganic N content, and we determined microbial communities with amplicon sequencing. With the remaining soil, we created our dilution inocula to perform the second part of our experiment.

### 2.3 Synthetic soil

#### 2.3.1 Synthetic soil preparation

To create synthetic soil, we adapted the protocol from Guenet et al. (2011). Briefly, we mixed 20 g of kaolinite and 10 g of bentonite with 300 mL of water for 2 days at 125 rpm. After two days, we sonicated the soil slurry at maximum speed for 10 min, using a Branson Sonifier 450 (Branson Ultrasonics Corporation, Danbury, CT, USA), and we centrifuged the mixture at 2000 x g for 20 min. We discarded the supernatant, added 70 g of sand and let the mixture dry at 4 °C for 3 days in a cooled chamber. After creating 3 kg of dried synthetic soil, we added 60 g of humic acid (Sigma-Aldrich; H16752) and autoclaved it. Humic acid is the only carbon source in our synthetic soil, containing only mineral material. We then split up the dried synthetic soil into 60 magenta boxes, and we added 5.5 mL of sulfuric acid (0.36 M) to obtain a pH around 5.5 – 6 (measured in a 1:2 soil to CaCL_2_, 0.01 M ratio), corresponding to the average pH of the two native soils.

#### 2.3.2 Microbial necromass preparation

We grew two microbes [i.e., *Bacillus cereus* (in Trypto Soy Broth) and *Penicillium sp.* (on Potato Dextrose Agar)] to create our microbial necromass. We centrifuged bacterial cells at 4500 rpm during 10 min and washed them with a ratio of 2 of cell culture to 1 citrate buffer (0.1 M, pH 7) to harvest them. We harvested fungal mycelia from plates by scraping them with a scalpel. After creating enough bacterial and fungal biomass, we autoclaved it twice for 60 min at 121 °C to create the necromass. We flash-freeze and lyophilized fungal and bacterial necromass before crushing it in liquid nitrogen using a mortar and pestle and adding it to the synthetic soil.

#### 2.3.3 Microbial dilution inoculation and necromass addition

After 9 weeks, we blended the equivalent of 10 g of dried soil from each magenta box (from the native soil experiment) for 2 min at maximum speed into 40 mL of water to create our synthetic soil inocula. We added 10 mL of each soil slurry into synthetic soil (0.05 g soil inoculum g^-1^ synthetic soil), mixed the synthetic soil and incubated it for two days before adding microbial necromass. For crop communities, we added 0.16 g of fungal necromass (C:N ratio = 9.72) and 0.08 g of bacterial necromass (C:N ratio = 3.84).

For forest communities, we added 0.69 g of fungal necromass and 0.29 g of bacterial necromass. We used these amounts to match the total amount of C, N and the C: N ratio of the native soil and the different amounts for both was calculated to match the amount in real conditions (Simpson et al., 2007) (Wang et al., 2021). According to these articles, there is more necromass in forest than crop soil and there is more fungal necromass than bacterial necromass.

### 2.4 Native and synthetic soil

#### 2.4.1 DNA extraction and sequencing

We extracted soil DNA from 0.25 g of soil samples using the DNeasy PowerLyzer PowerSoil Kit (Qiagen) following the manufacturer’s protocol. DNA was eluted in 25 μL of elution buffer and was stored at −20 °C until further analyses. We prepared libraries for 16S and ITS amplicon sequencing according to Illumina’s 16S Metagenomic Sequencing Library Preparation guide (Part#15044223Rev.B). We prepared libraries for the bacterial 16S rRNA gene using primers 16S: 515Y and 926R, and for the fungal ITS1-ITS2 region, we used the primers ITS9F and ITS4R. NextSeq (Illumina) sequencing (PE300 10M reads) was performed at the Centre d’expertise et de service Génome Québec (CESGQ, Montréal, Canada).

#### 2.4.2 Enzymatic potential activities quantification

We quantified the potential activity of three exoenzymes involved in SOM depolymerization (proteases, deaminases and β-glucosidases) using colorimetric methods. Every protocol follows the same principle: soil samples were incubated with an excess of substrate under optimal conditions. Therefore, the result is the potential activity, meaning that they do not reflect real-time activity in the soil. We quantified potential activities according to protocols previously described in L’Espérance et al. (2025). Briefly, we incubated 1 g of dried sieved (2 mm) soil with 2.5 mL sodium caseinate (2%) and 2.5 mL Tris buffer for two hours at 50 °C and agitated at 100 rpm. We added 5 mL Trichloroacetic acid (1 g L^-1^), 0.75 mL Na_2_CO_3_ and 0.25 mL Folin & Ciocalteu’s phenol (1:3). We measured the absorbance at 650 -700 nm. Potential protease activities are expressed in μmol tyrosine released g^−1^ dry soil h^−1^ incubation and was quantified using a tyrosine standard curve (2-fold serial dilution from 1.0 to 0.005 mM). To quantify deaminase potential activities, we incubated 1 g of dried sieved (2 mm) soil with 1.2-diamino-4-nitrobenzene (0.06 g L^-1^) for 20 h at 25 °C, 100 rpm. We then extracted twice the remaining substrate using methanol 100 % and filtered (Whatman no1) the solution. We measured the absorbance at 405 nm. Potential activities were quantified using a 1.2-diamino-4-nitrobenzene standard curve (0/ 0.008/ 0.011/ 0.014/ 0.019/ 0.025/ 0.034/ 0.045/ 0.06 g L^-1^ of diamino-4-nitrobenzene). Finally, to quantify β-glucosidase potential activity, we incubated 1 g of dried sieved (2 mm) soil with a buffer containing p-nitrophenyl glycoside (25 mM) at 37 °C for 1 h. We then added calcium chloride (0.5 M) and Tris buffer (0.1 M, pH 12). We measured absorbance at 415 nm. Potential activities are expressed in μmol p-nitrophenol released g^−1^ dry soil h^−1^ incubation and was quantified using a p-nitrophenol standard curve (2-fold serial dilution from 0.5 to 0.0078 g L^-1^).

#### 2.4.3 Available inorganic nitrogen quantification

We extracted inorganic N from the soil by incubating 1 g of dried sieved soil in 10 mL of KCl (2 M) for 1 h at 25 °C 100 rpm. After incubation, we filtered (Whatman no. 42) the soil slurry and the filtered soil extracts were kept at 4 °C until further analyses. We quantified ammonium and nitrate in our samples as previously described (L’Espérance et al., 2025). Briefly, we used 100 μL of the sample and added 50 μL of citrate reagent, 50 μL of phenylphenol nitroprusside, 25 μL of buffered hypochlorite reagent and 50 μL of deionized water to quantify ammonium content. We incubated 45 min at room temperature, and we measured the absorbance at 660 nm. Ammonium concentration is expressed in mg g^-1^ of air-dried soil and quantified using a (NH_4_)_2_SO_4_ standard curve (2-fold serial dilution from 20 to 0.0195 mg L^-1^). To quantify nitrate, we used 100 μL of the sample and added 100 μL of VCl_3_ and 100 μL of the Griess reagent mix (1:1 Griess solution 1 and 2). We incubated the solution for 1 h at 37 °C and we measured absorbance at 540 nm. Nitrate concentration is expressed in mg g^-1^ of soil and quantified with a standard curve of KNO_3_ (2-fold serial dilution from 10 to 0.0195 mg L^-1^). For the synthetic soil experiment, we assessed mineralization rate by quantifying ammonification rate (ammonium content after 14 days minus initial ammonium content, expressed in mg of N kg^-1^ day^-1^) and nitrification rate (nitrate content after 14 days minus initial nitrate content, expressed in mg of N kg^-1^ day^-1^).

#### 2.4.4 Bioinformatic analyses

We analyzed amplicon sequencing data (16S rRNA gene and ITS region) using the DADA2 pipeline (v.1.38.0) in R (v.4.5.1) to process raw reads (Callahan et al., 2016). For 16S, we used truncLen (280,250) and maxEE (2,2) to filter and trim forward and reverse reads. We used the pooled method for removing chimeras, and we assigned taxonomy with the SILVA reference database (v.138.1). After the assignation, we removed chloroplasts and mitochondria from our dataset before further analysis. For ITS, we used truncLen (270,250) and maxEE (2,2) to filter and trim forward and reverse reads. We used the pooled method for removing chimeras, and we assigned taxonomy with the UNITE reference database (v.8.3).

#### 2.4.5 Statistical analysis and data visualization

We performed statistical analyses in R (version 4.5.1, *The R Foundation for Statistical Computing*) and created figures using the ggplot2 package (v.4.0.2). For testing the effect of the microbial dilution on potential enzymatic activities, soil nitrogen content, and mineralization rate, we performed analysis of variance (ANOVA) when the assumptions of normal distribution of residuals and variance homogeneity were met. When the assumptions were met and the analysis of variance (ANOVA) was statistically significant (p< 0.05), we performed Tukey multiple comparisons to reveal significant differences between groups. If these assumptions were not met, we performed a Kruskal-Wallis non-parametric test. We performed Kruskal-Wallis non-parametric tests to determine the effect of microbial dilution on alpha-diversity and relative abundance data. To analyze the bacterial community structure, we first transformed our dataset using the Hellinger transformation. We then used Euclidean distance matrices based on our transformed data to visualize the community structure using a principal coordinate analysis (PCoA). We tested the effect of microbial dilution on the community using PERMANOVA (1000 permutations). We then removed our control and used the 10 first PCoA axis in linear models to explain each exoenzymes potential activities and mineralization processes. When the model was significant, we performed Spearman correlation to find positively correlated ASVs. Briefly, positive correlation means that when ASVs abundances increase, the responses variable increase (exoenzymes potential activities or mineralization processes).

#### 2.4.6 Data availability

The R code used to analyze the data and generate the figures is available on GitHub (https://github.com/le-labo-yergeau/). Raw sequencing reads were deposited in the NCBI SRA repository under BioProject accession PRJNA1468094.

## 3. Results

### 3.1 Native soil

We successfully altered the alpha and beta diversity of bacterial and fungal communities using the dilution technique across both soil types (Table S1, Fig. 1 and Fig. 2). As expected, the D0 treatment has the highest Shannon index and the D8 the lowest, except for the forest bacterial communities (Fig. 1.A; 1.D and Fig. 2.A; 2.D). Specifically, for crop bacterial communities, D6, D8 and CTL are statistically similar (Fig. 1.A). For forest bacterial communities, D4, D6, D8 and CTL are statistically similar (Fig. 1.D) For crop fungal communities, there is no significant difference, even if D0 has the highest Shannon index and D8 the lowest (Fig. 2.A). Finally, for forest fungal communities, D6, D8 and CTL are statistically similar (Fig. 2.D). Our controls (sterilized soil without any addition of microbes) have a highly variable Shannon diversity index (CV = 58.96, 61.89, 92.84 and 105.33 % in Fig. 1.A and 1.D and Fig. 2.A and 2.D respectively), suggesting that some microbes survived the sterilization process in some of the replicates. There is a clear separation of bacterial community composition by dilution treatments. Dilution D0 and D2 cluster together and are separate from the D6 - D8 cluster (Fig. 1B; 1.E). Fungal community patterns are similar but less pronounced than those of bacterial communities. In both soils, there is a separation of D6, D8 and CTL, and the clustering of D0, D2 and D4 (Fig. 2B; 2E).

**Figure 1:**
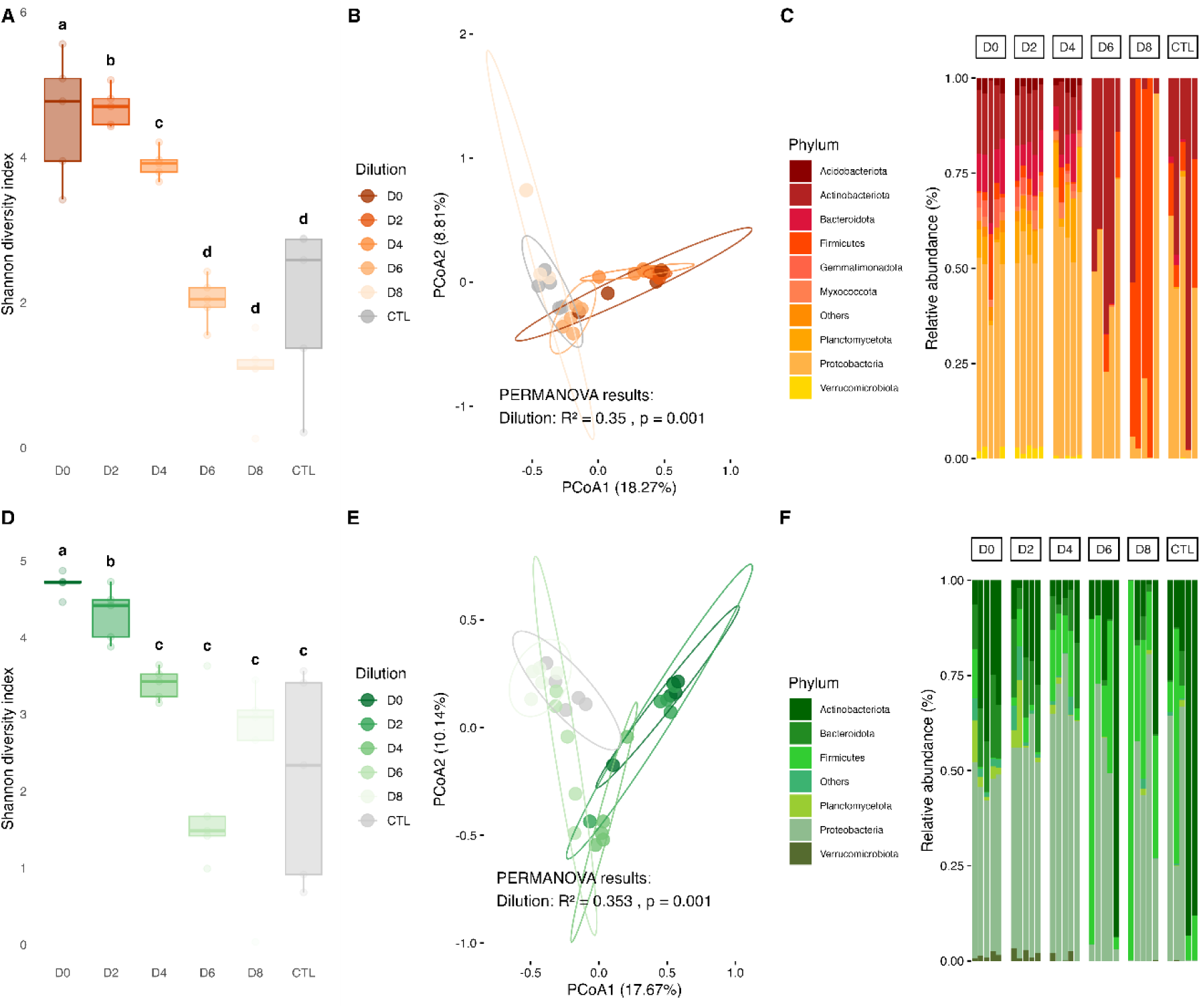
Alpha and Beta diversity of bacterial (16S) communities from diluted communities from crop soil (A, B, C) and forest soil (D, E, F) in their native soil after 9 weeks of incubation. Alpha diversity (A, D) is represented by Shannon diversity indexes, and significant results from Kruskal-Wallis and pairwise Wilcox tests adjusted with Benjamini-Hochberg correction correspond to letters above the boxplot (p<0.05). The boxplot represents the median (middle line), boxes represent the first and third quartiles and whiskers shows the extends of the data, while individual dots represent outliers. Beta diversity (B, E) is represented using a Euclidean distance matrix and principal coordinate analysis. The R2 and p-value (B, E) correspond to the PERMANOVA results for the dilution treatment.

**Figure 2:**
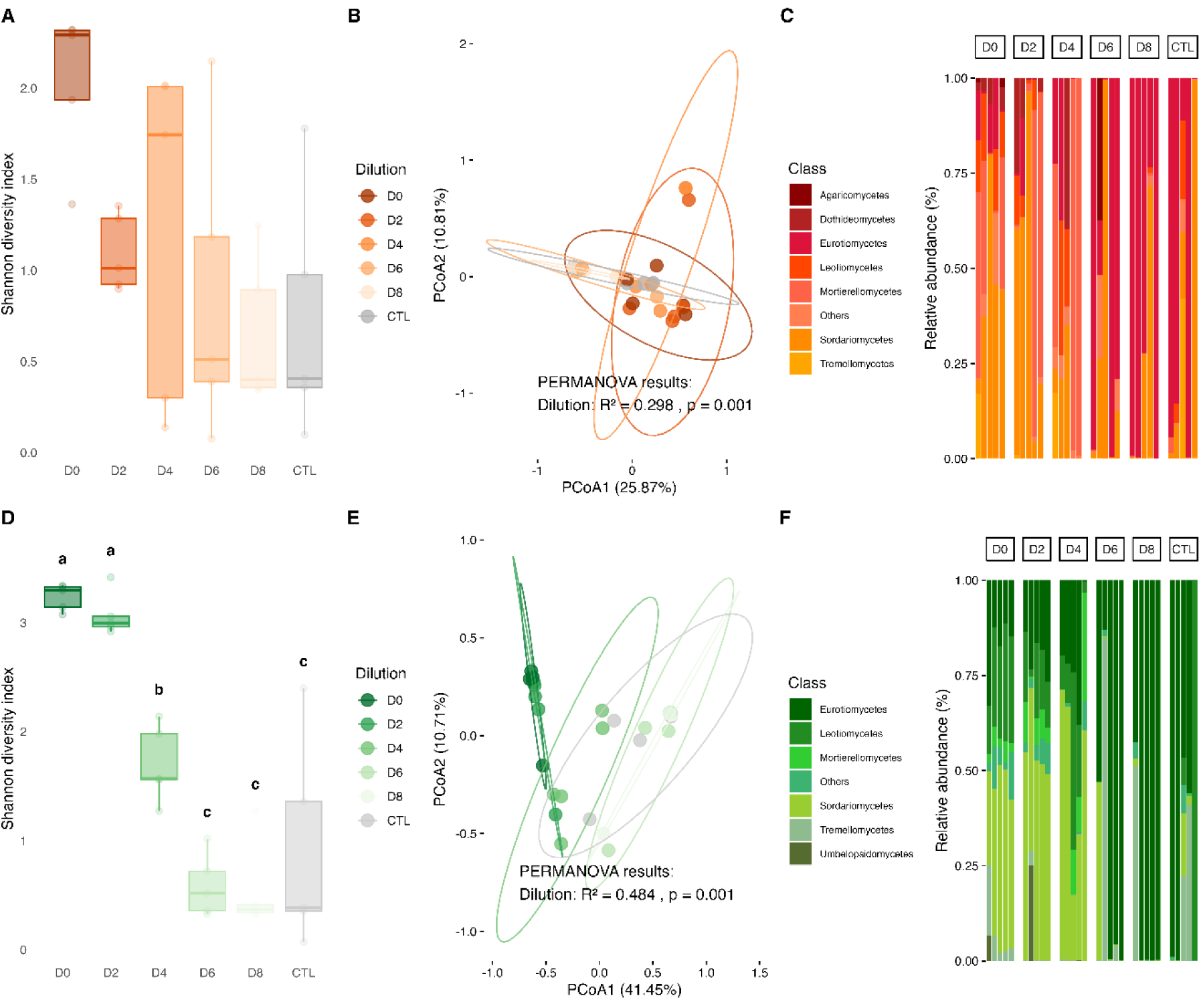
Alpha and Beta diversity of fungal (ITS) communities from diluted communities from crop soil (A, B, C) and forest soil (D, E, F) in their native soil after 9 weeks of incubation. Alpha diversity (A, D) is represented by Shannon diversity indexes, and significant results from Kruskal-Wallis and pairwise Wilcox tests adjusted with Benjamini-Hochberg correction correspond to letters above the boxplot (p<0.05). The boxplot represents the median (middle line), boxes represent the first and third quartiles and whiskers shows the extends of the data, while individual dots represent outliers. Beta diversity (B, E) is represented using a Euclidean distance matrix and principal coordinate analysis. The R2 and p-value (B, E) correspond to the PERMANOVA results for the dilution treatment.

Bacterial community composition varied significantly among dilution in their native soil (crop soil p-value= 0.009 and forest soil p-value= 0.049) (Fig. 1.C; 1.F; 3.A; 3.B). Bacterial communities from the crop soil were more variable at dilution D6 and D8 than less diluted communities (D2 and D4) (Fig. 1.C; 3.A). The variation in bacterial communities from the forest soil increased with dilution, but pairwise Wilcoxon test did not reveal significant differences between specific dilution pairs (Fig. 3.B). On the other hand, fungal community composition did not vary significantly among dilution in their native soil (crop soil p-value=0.429 and forest soil p-value=0.197) (Fig. 2.C; 2.F; 3.C; 3.D).

**Figure 3:**
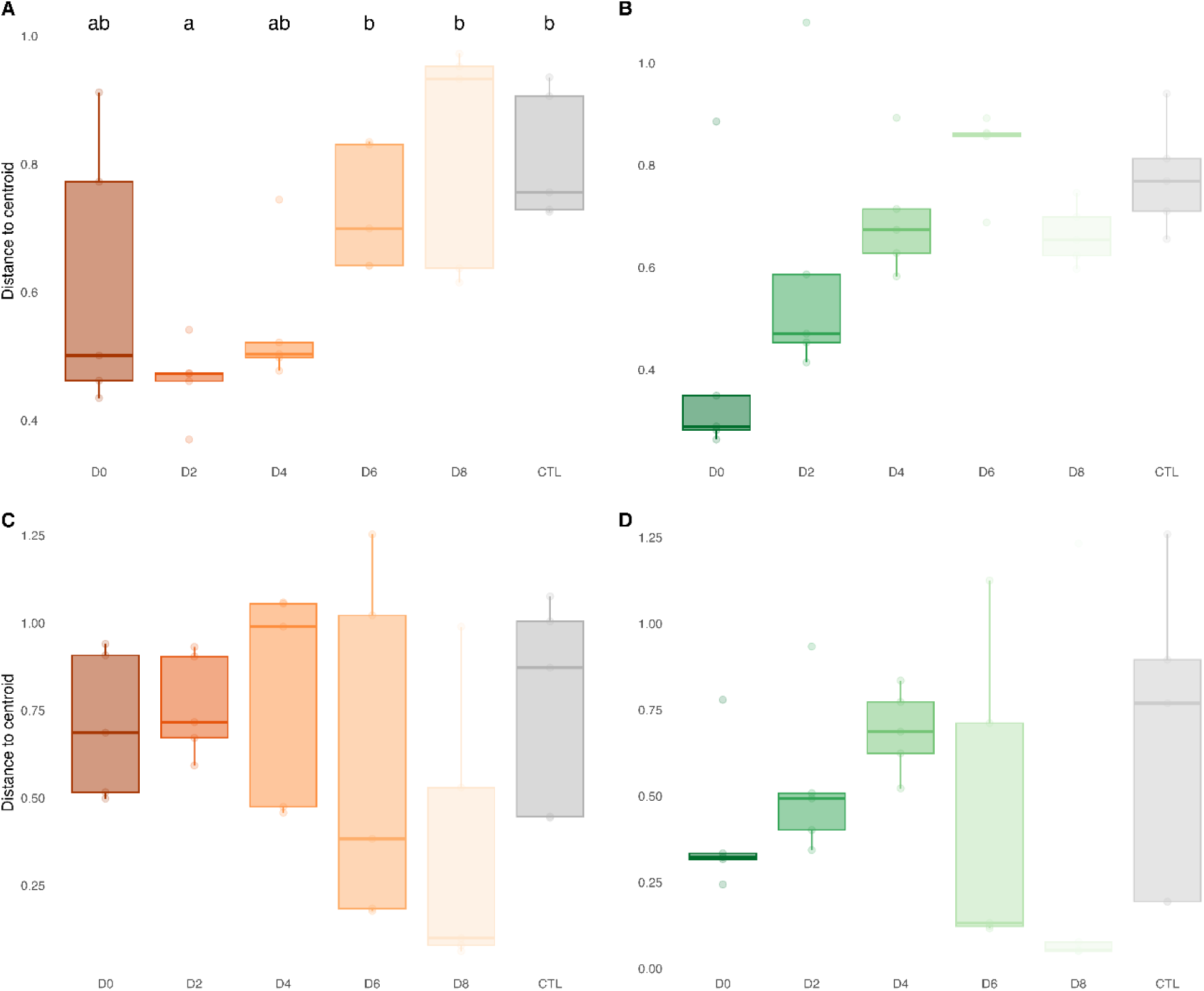
Beta dispersion of bacterial (A, B) and fungal (C, D) communities from diluted communities from crop soil (A, C) and forest soil (B, D) in their native soil after 9 weeks of incubation. Significant results from Kruskal-Wallis and pairwise Wilcox tests adjusted with Benjamini-Hochberg correction, correspond to letters above the boxplot (p<0.05). The boxplot represents the median (middle line), boxes represent the first and third quartiles and whiskers shows the extends of the data, while individual dots represent outliers.

After 9 weeks of incubation in their native soil, dilution treatments did not have a significant effect on protease, β-glucosidase and deaminase potential activities of crop soil microbial communities (Fig. 4A, 4B and 4C). For forest soil microbial communities, there was a significant effect of dilution treatments on β-glucosidase potential activity (p-value = 0.0396), but Tukey multiple comparisons of means could not single out the treatments responsible for this significant difference (Table 1 and Fig. 4.E). We can still observe tendencies towards higher potential β-glucosidase activities in intermediate dilution treatments (D4 and D6). We did not find any significant correlation between Shannon indexes and exoenzymes potential activities (proteases, β-glucosidase and deaminase potential activities, data not shown).

**Figure 4:**
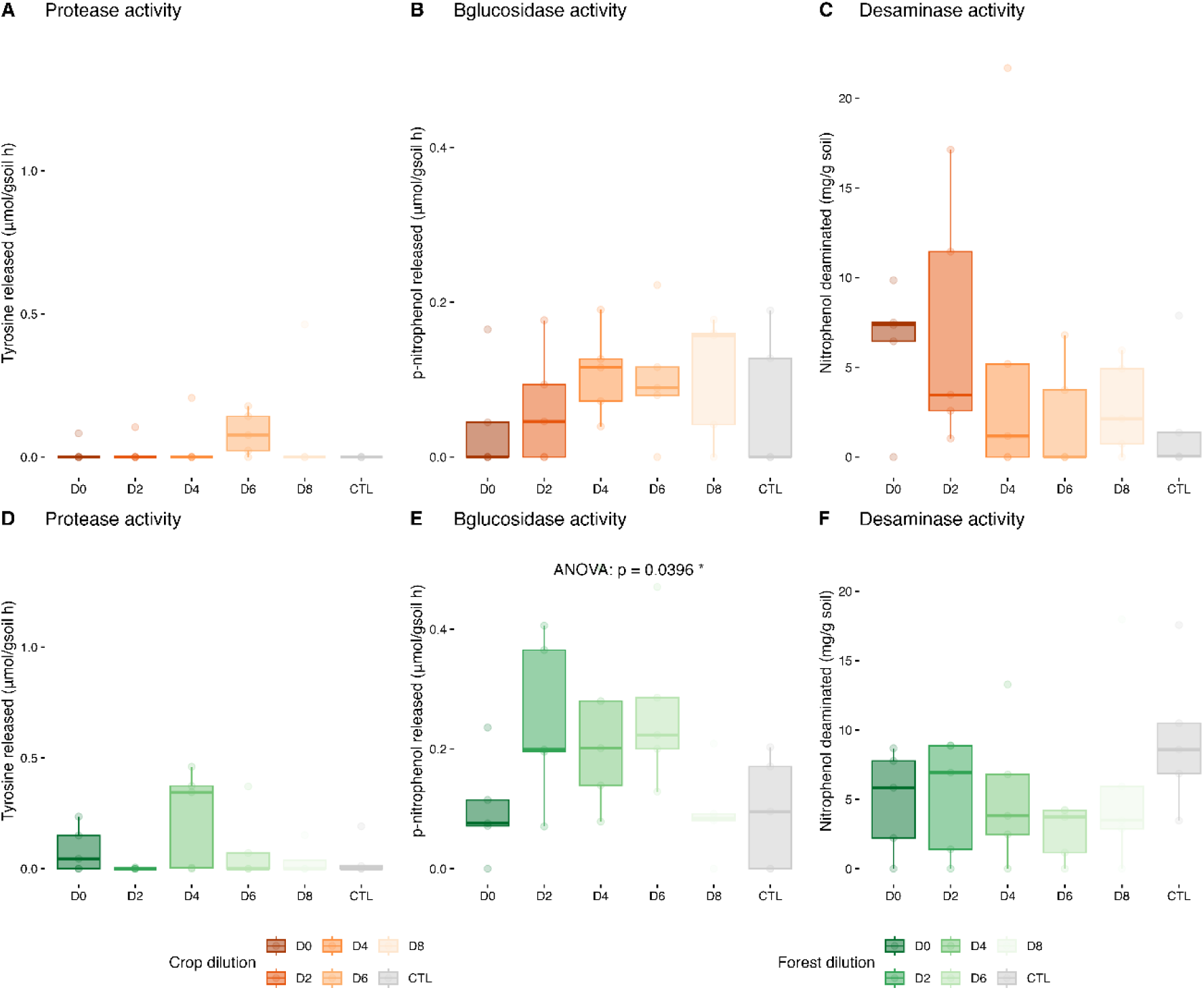
Protease, β-glucosidase and deaminases’ potential activities of diluted communities after 9 weeks of incubation in crop soil (A, B, C) and forest soil (D, E, F). Effects of dilution treatments are represented. Significant results from Kruskal-Wallis or ANOVA followed by Tukey multiple comparisons for each enzyme’s potential activities are identified with a *(p<0.05). The boxplot represents the median (middle line), boxes represent the first and third quartiles and whiskers shows the extends of the data, while individual dots represent outliers.

**Table 1:** Effects of dilution treatment on proteases, β-glucosidases and deaminases’ potential activities and ammonium and nitrate content in the native soil.

| Soil | Analysis | Response variable | Factor | Statistic | df | p-value |
| --- | --- | --- | --- | --- | --- | --- |
| Crop | Kruskal | Protease activity | Dilution | 7.567 | 5 | 0.182 |
| | ANOVA | β-glucosidase activity | Dilution | 1.102 | 5 | 0.390 |
|  | ANOVA | Deaminase activity | Dilution | 1.362 | 5 | 0.280 |
|  | Kruskal | Ammonium content | Dilution | 3.708 | 5 | 0.592 |
|  | ANOVA | **Nitrate content** | **Dilution** | **2.242** | **5** | **0.090** |
| Forest | Kruskal | Protease activity | Dilution | 5.326 | 5 | 0.377 |
|  | ANOVA | **β-glucosidase activity** | **Dilution** | **2.903** | **5** | **0.040** * |
|  | ANOVA | Deaminase activity | Dilution | 1.014 | 5 | 0.435 |
|  | Kruskal | **Ammonium content** | **Dilution** | **17.516** | **5** | **0.004** ** |
|  | Kruskal | Nitrate content | Dilution | 2.396 | 5 | 0.792 |

. : p<0.1; \*: p<0.05; \*\*p<0.01; \*\*\*:p<0.001

After 9 weeks of incubation, there was significantly more ammonium in the forest soil dilutions D4 and D6 than in the non-diluted sample (p-value = 0.004) (Table 1 and Fig. 5.C). Specifically, there is 11 time more ammonium in the dilution D4 and 9 time more ammonium in the dilution D6 than in the non-diluted sample. Ammonium content in the crop soil was not significantly different between dilution treatments. There was a trend toward more nitrate in the non-diluted sample from crop soil than in the other dilutions, (p-value = 0.090). There was no significant difference in nitrate content among dilutions treatments in the forest soil.

**Figure 5:**
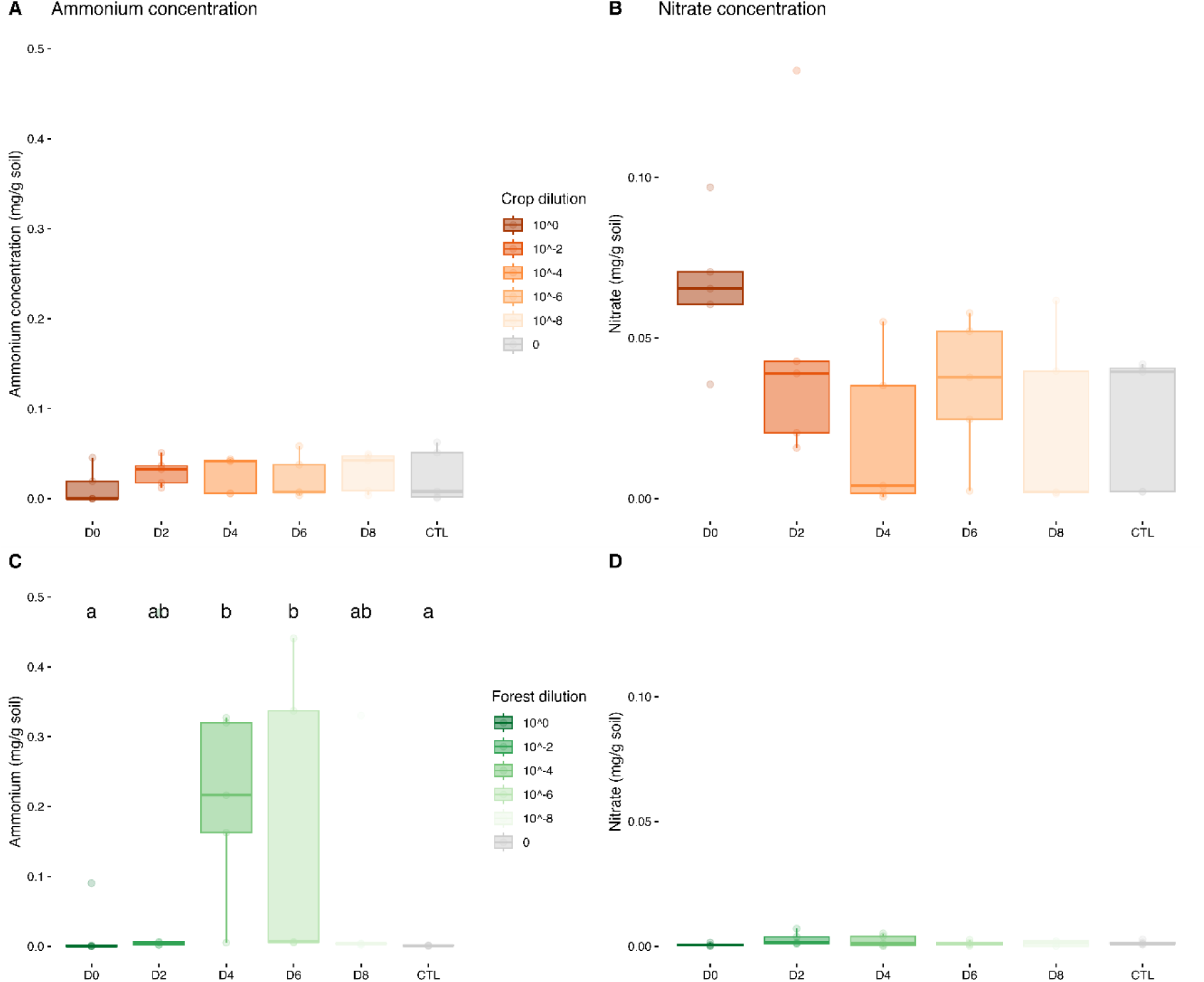
Ammonium and nitrate concentration in crop soil (A, B) and forest soil (C, D) containing diluted communities after 9 weeks of incubation. Effects of dilution treatments are represented. Significant results from Kruskal-Wallis or ANOVA followed by Tukey multiple comparisons correspond to letters above the boxplot (p<0.05). The boxplot represents the median (middle line), boxes represent the first and third quartiles and whiskers shows the extends of the data, while individual dots represent outliers.

### 3.2 Synthetic soil

After 2 weeks of incubation with microbial necromass in the synthetic soil, the alpha and beta diversity of bacterial and fungal communities were distinct between dilution treatments (Table S2, Fig. 6 and Fig.7). Specifically, for crop bacterial communities, D0 and D2 has the highest Shannon index and D6, D8 and CTL are statistically similar and had the lowest Shannon index (Fig. 6.A). For forest bacterial communities, D0 and D2 had the highest Shannon index and D4, D6, D8 and CTL are statistically similar and had the lowest Shannon index (Fig. 6.D) For crop fungal communities, D4 was statistically similar to D0 and D2 and has the highest Shannon index and D6 and D8 had the lowest Shannon index (Fig. 7.A). Finally, for forest fungal communities, D2 had the highest Shannon index and was statistically similar to D0 and D4, whereas D6 and D8 had the lowest Shannon index (Fig. 7.D). Our controls (sterilized soil without any addition of microbes) were variable, suggesting that some microbes survived the sterilization process and were able to colonize the synthetic soil after the transfer (CV = 19.54, 27.56, 119.49 and 98.43 % in Fig. 6.A and 6.D and Fig. 7.A and 7.D respectively). Bacterial community composition (Fig. 6.B; 6.E) varied between replicates and this variability was more pronounced in the synthetic soil that received the more diluted treatment (D6 and D8) and the controls. Fungal community composition also varied between replicates, but the pattern was less clear than for bacteria (Fig. 7.B; 7.E). Interestingly, there is a clear separation in the composition of both fungal and bacterial communities by dilution treatment. The less diluted treatments (D0, D2, D4) created tight clusters, especially for bacterial communities (Fig. 6.B; 6.E; 7.B; 7.E).

**Figure 6:**
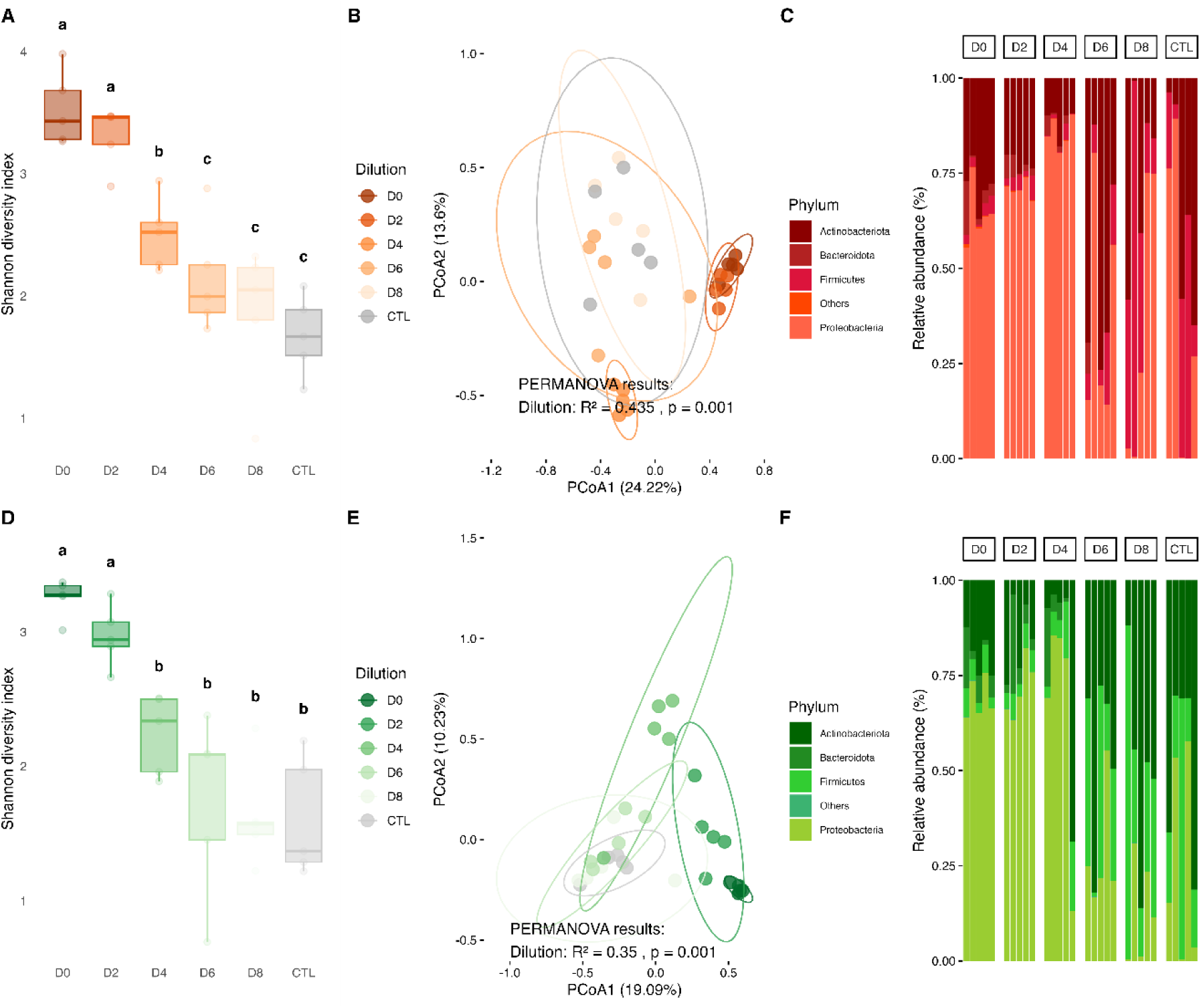
Alpha and Beta diversity of bacterial (16S) communities from diluted communities from crop soil (A, B, C) and forest soil (D, E, F) in the synthetic soil with microbial necromass after 2 weeks of incubation. Alpha diversity (A, D) is represented by Shannon diversity indexes, and significant results from Kruskal-Wallis and pairwise Wilcox tests correspond to letters above the boxplot (p<0.05). The boxplot represents the median (middle line), boxes represent the first and third quartiles and whiskers shows the extends of the data, while individual dots represent outliers. Beta diversity (B, E) is represented using a Euclidean distance matrix and principal coordinate analysis. The R2 and p-value correspond to the PERMANOVA results for the dilution treatment.

**Figure 7:**
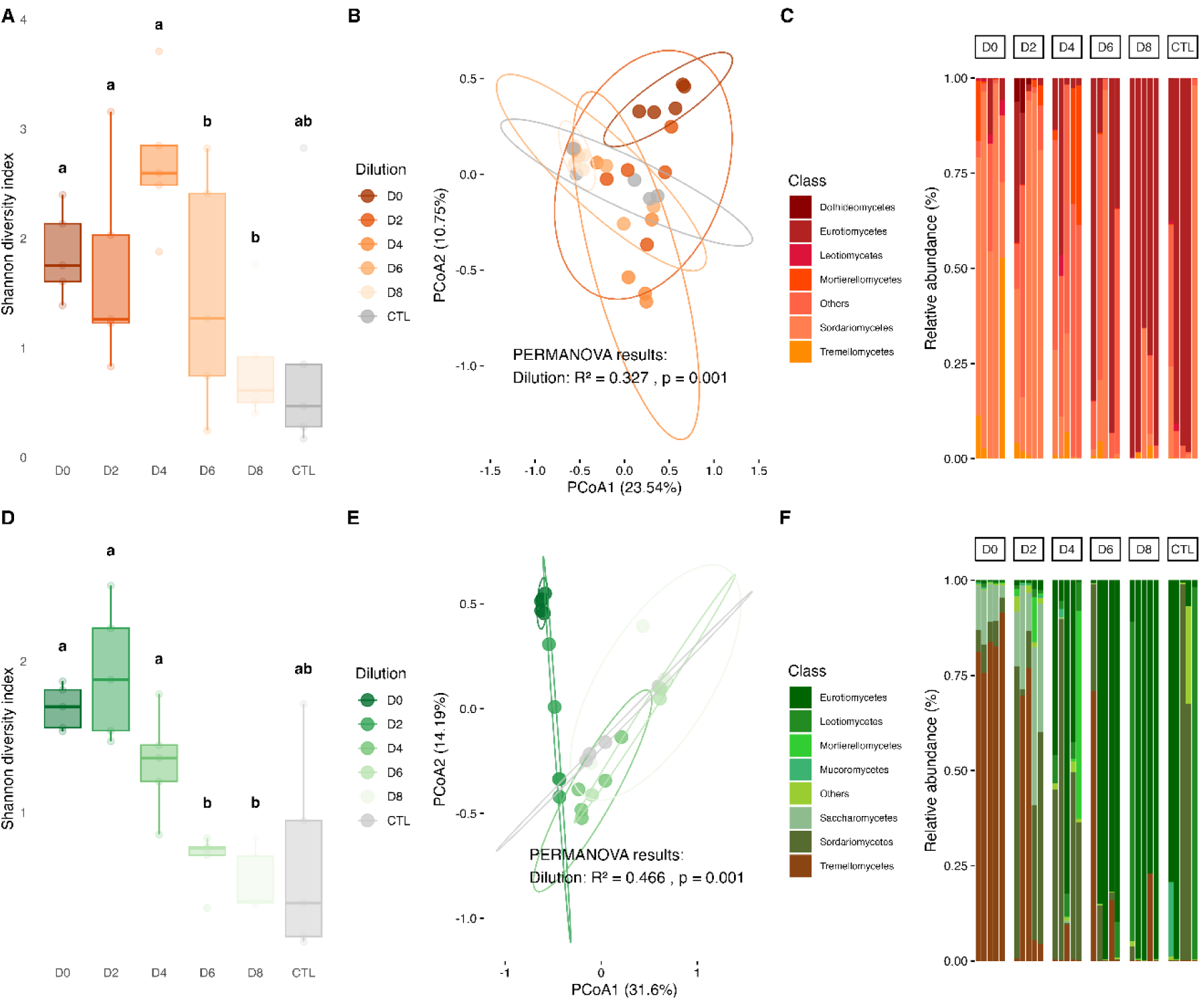
Alpha and Beta diversity of fungal (ITS) communities from diluted communities from crop soil (A, B, C) and forest soil (D, E, F) in the synthetic soil with microbial necromass after 2 weeks of incubation. Alpha diversity (A, D) is represented by Shannon diversity indexes, and significant results from Kruskal-Wallis and pairwise Wilcox tests correspond to letters above the boxplot (p<0.05). The boxplot represents the median (middle line), boxes represent the first and third quartiles and whiskers shows the extends of the data, while individual dots represent outliers. Beta diversity (B, E) is represented using a Euclidean distance matrix and principal coordinate analysis. The R2 and p-value correspond to the PERMANOVA results for the dilution treatment.

The variability of the bacterial community composition changed significantly among dilutions in the synthetic soil (crop soil p-value=0.002 and forest soil p-value=0.009) (Fig. 6C; 6F; 8A; 8B). Bacterial communities from crop and forest soils were more variable at dilution D6 and D8 than less diluted communities (Fig. 8.A; 8.B). Fungal communities from the forest soil varied among dilutions in the synthetic soil (p-value = 0.033), but pairwise Wilcoxon tests did not reveal significant differences between specific dilution (Fig. 8.C; 8.D).

**Figure 8:**
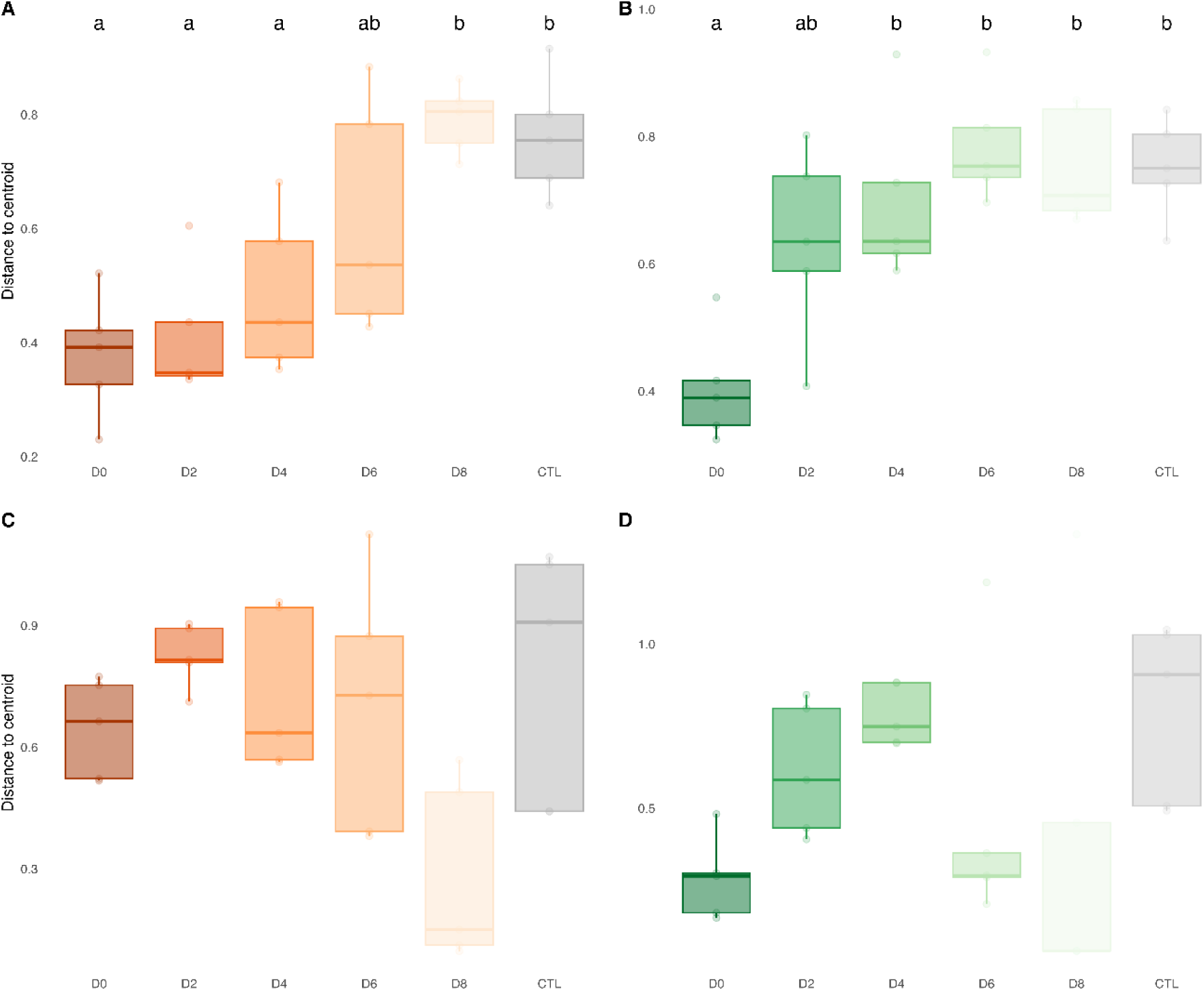
Beta dispersion of bacterial (A, B) and fungal (C, D) communities from diluted communities from crop soil (A, C) and forest soil (B, D) in the synthetic soil after 2 weeks of incubation with microbial necromass. Significant results from Kruskal-Wallis and pairwise Wilcox tests adjusted with Benjamini-Hochberg correction, correspond to letters above the boxplot (p<0.05).

Two weeks after microbial necromass addition, dilution treatment had a significant effect on β-glucosidase potential activities of crop soil communities (p-value = 0.025) and protease potential activities of forest soil communities (p-value = 0.040) (Table 3 and Fig. 9). β-glucosidase potential activities were significantly different between the crop soil microbial communities D4 and D2. Specifically, D4 microbial communities had about 8 times higher β-glucosidase potential activities than D2 communities. β-glucosidase potential activities were different and nearly significant (p-value = 0.079) among forest soil microbial communities. Dilution treatments of the forest soil microbial communities significantly influenced protease potential activities, but the Tukey multiple comparisons of means failed to reveal which groups were significantly different. Higher protease potential activities were found in the undiluted communities (i.e., D0) and D4 and D6, approximately 5 times higher than D2 and D8. We did not find any significant correlation between Shannon indexes and exoenzymes potential activities (proteases, β-glucosidase and deaminase potential activities) and mineralization processes (ammonification and nitrification).

**Figure 9:**
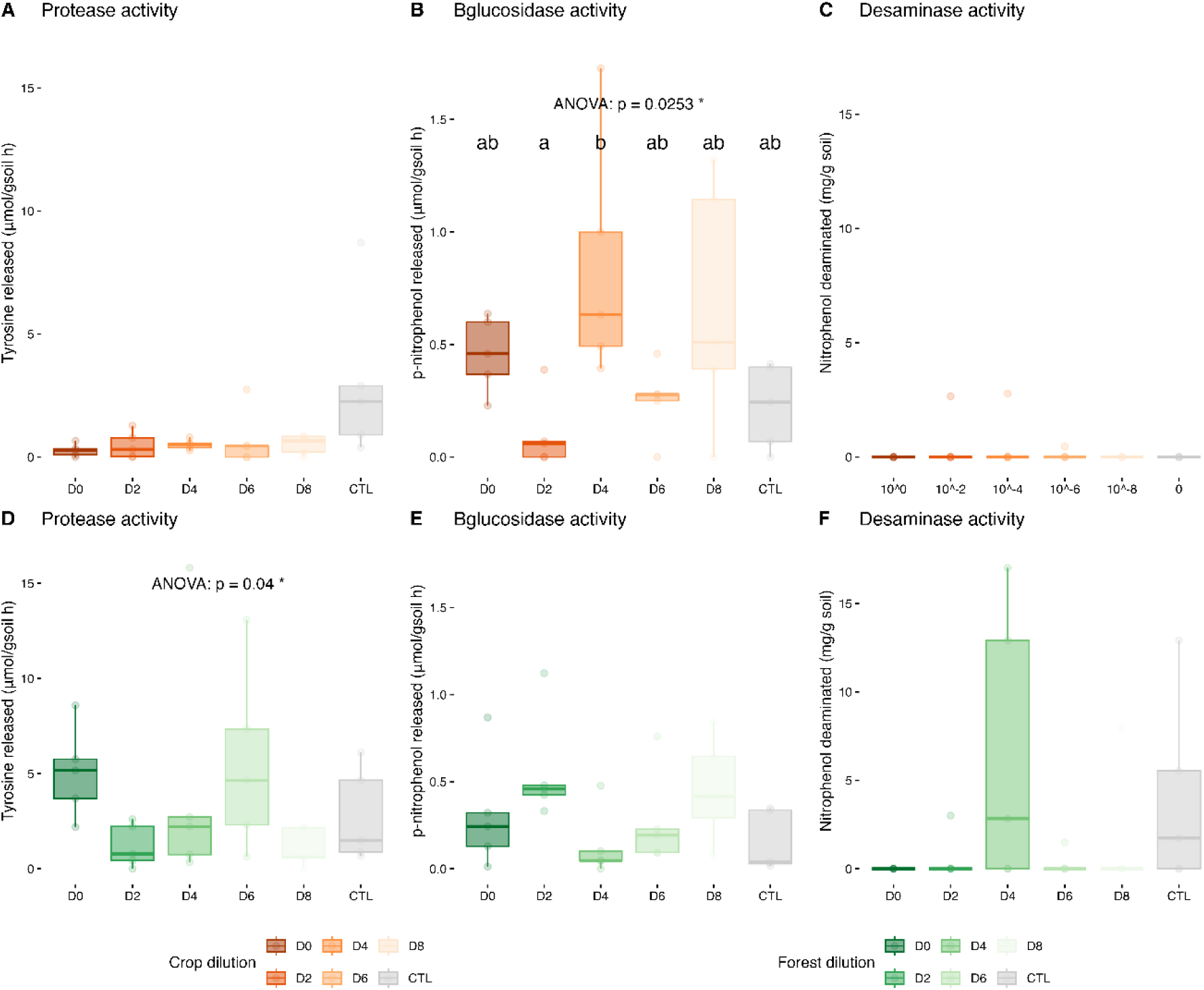
Protease, β-glucosidase and deaminases’ potential activities of diluted communities from crop soil (A, B, C) and forest soil (D, E, F) after 2 weeks of incubation in synthetic soil with microbial necromass. Effects of dilution treatments are represented. Significant results from Kruskal-Wallis or ANOVA followed by Tukey multiple comparisons for each enzyme’s potential activities (see Table 1) are identified with a *(p<0.05). The boxplot represents the median (middle line), boxes represent the first and third quartiles and whiskers shows the extends of the data, while individual dots represent outliers.

**Table 3:** Effects of dilution treatment on proteases, β-glucosidases and deaminases potential activities and ammonium and nitrate content in the synthetic soil.

| Soil | Analysis | Response variable | Factor | Statistic | df | p-value |
| --- | --- | --- | --- | --- | --- | --- |
| Crop | Kruskal | Protease activity | Dilution | 8.366 | 5 | 0.137 |
|  | ANOVA | **β-glucosidase activity** | **Dilution** | **3.279** | **5** | **0.025 \*** |
|  | Kruskal | Deaminase activity | Dilution | 3.231 | 5 | 0.665 |
|  | ANOVA | Ammonium content | Dilution | 0.469 | 5 | 0.795 |
|  | ANOVA | Nitrate content | Dilution | 0.375 | 5 | 0.860 |
| Forest | ANOVA | **Protease activity** | **Dilution** | **2.893** | **5** | **0.0400 \*** |
|  | ANOVA | **β-glucosidase activity** | **Dilution** | **2.349** | **5** | **0.079 .** |
|  | Kruskal | Deaminase activity | Dilution | 7.739 | 5 | 0.171 |
|  | ANOVA | Ammonium content | Dilution | 1.083 | 5 | 0.400 |
|  | ANOVA | Nitrate content | Dilution | 0.460 | 5 | 0.801 |

. : p<0.1; \*: p<0.05; \*\*p<0.01; \*\*\*:p<0.001

Two weeks after microbial necromass addition in the synthetic soil, diluted microbial communities from crop and forest soil did not show a significant difference in mineralization rate (Table 4 and Fig. 10). When we separated mineralization into ammonification and nitrification, there was a difference in nitrification rate among diluted crop soil microbial communities (p-value = 0.012). D4 and D6 have a significantly lower nitrification rate than the control (about 68% lower).

**Figure 10:**
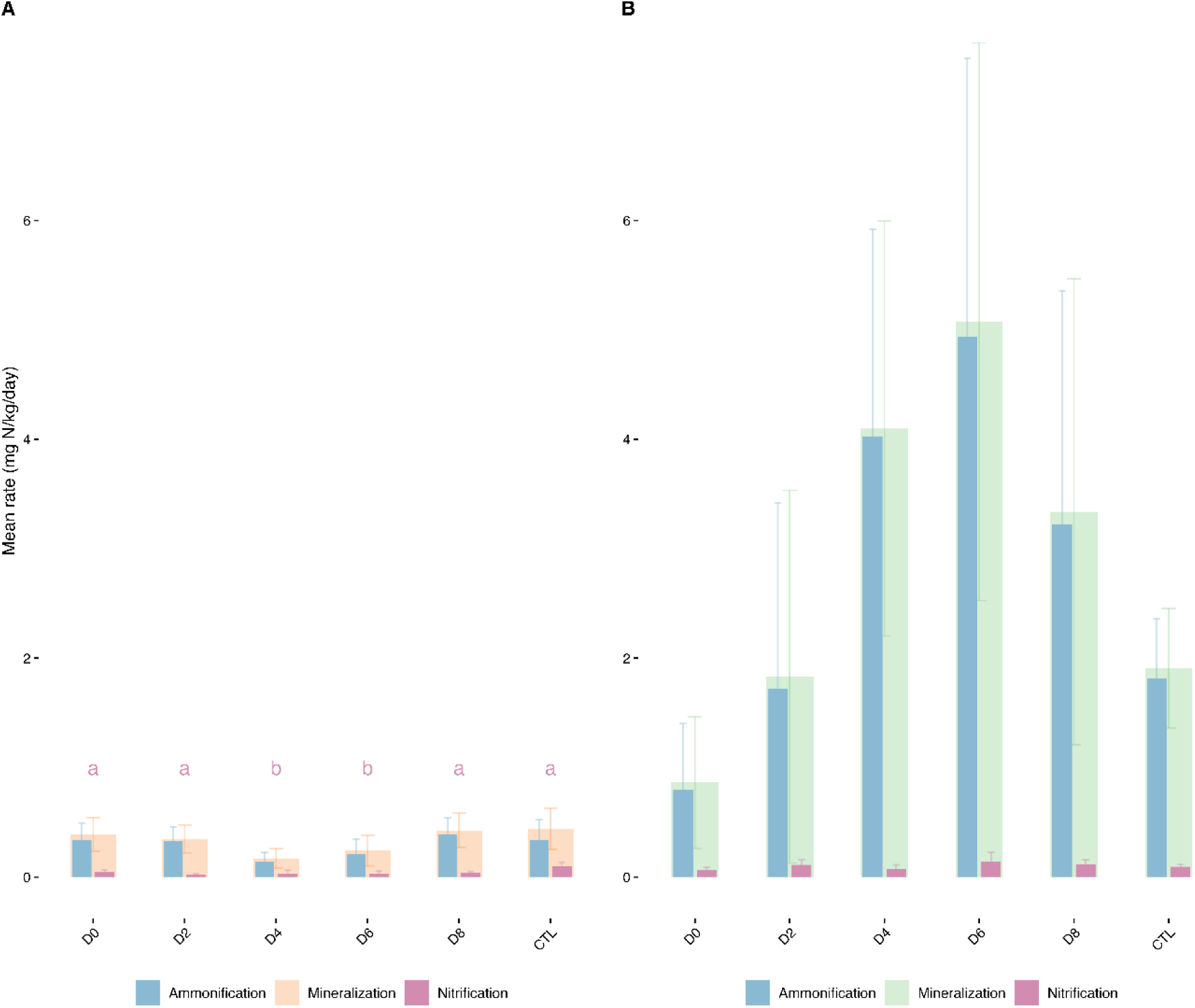
Mineralization rate (A: Crop soil microbial communities, B: Forest soil of the microbial necromass) after 2 weeks of incubation in the synthetic soil with microbial necromass. Significant results from ANOVA followed by Tukey multiple comparisons for each function are identified with a *(p<0.05).

**Table 4:** Effects of dilution treatment on mineralization, ammonification and nitrification rate in the synthetic soil analysed with ANOVA.

| Soil | Response variable | Factor | Statistic | df | p-value |
| --- | --- | --- | --- | --- | --- |
| Crop | Mineralization | Dilution | 1.337 | 5 | 0.289 |
|  | Ammonification | Dilution | 0.606 | 5 | 0.696 |
|  | **Nitrification** | **Dilution** | **3.931** | **5** | **0.012 \*** |
| Forest | Mineralization | Dilution | 1.052 | 5 | 0.425 |
|  | Ammonification | Dilution | 0.969 | 5 | 0.460 |
|  | Nitrification | Dilution | 0.012 | 5 | 0.882 |

. : p<0.1; \*: p<0.05; \*\*p<0.01; \*\*\*:p<0.001

We tested the effect of microbial community structure (PcoA axes) on protease, β-glucosidase and deaminase potential activities and ammonification and nitrification using linear models (Table 5). The microbial community structure from crop soil in the synthetic soil explained significantly enzymatic potential activities. Specifically, bacterial communities’ structure explained β-glucosidase activities (p-value = 0.044) and fungal communities structure explained protease activities (p-value = 0.019). The microbial community structure from forest soil also explained significantly enzymatic potential activities, and mineralization processes. Bacterial community’s structure explained protease (p-value = 0.003) and deaminase (p-value = 0.027) activities and both of mineralization processes (ammonification p-value = 0.0001 and nitrification p-value = 0.023). Fungal community’s structure explained deaminase activities (p-value = 0.021) and ammonification (p-value = 0.0002).

**Table 5:**
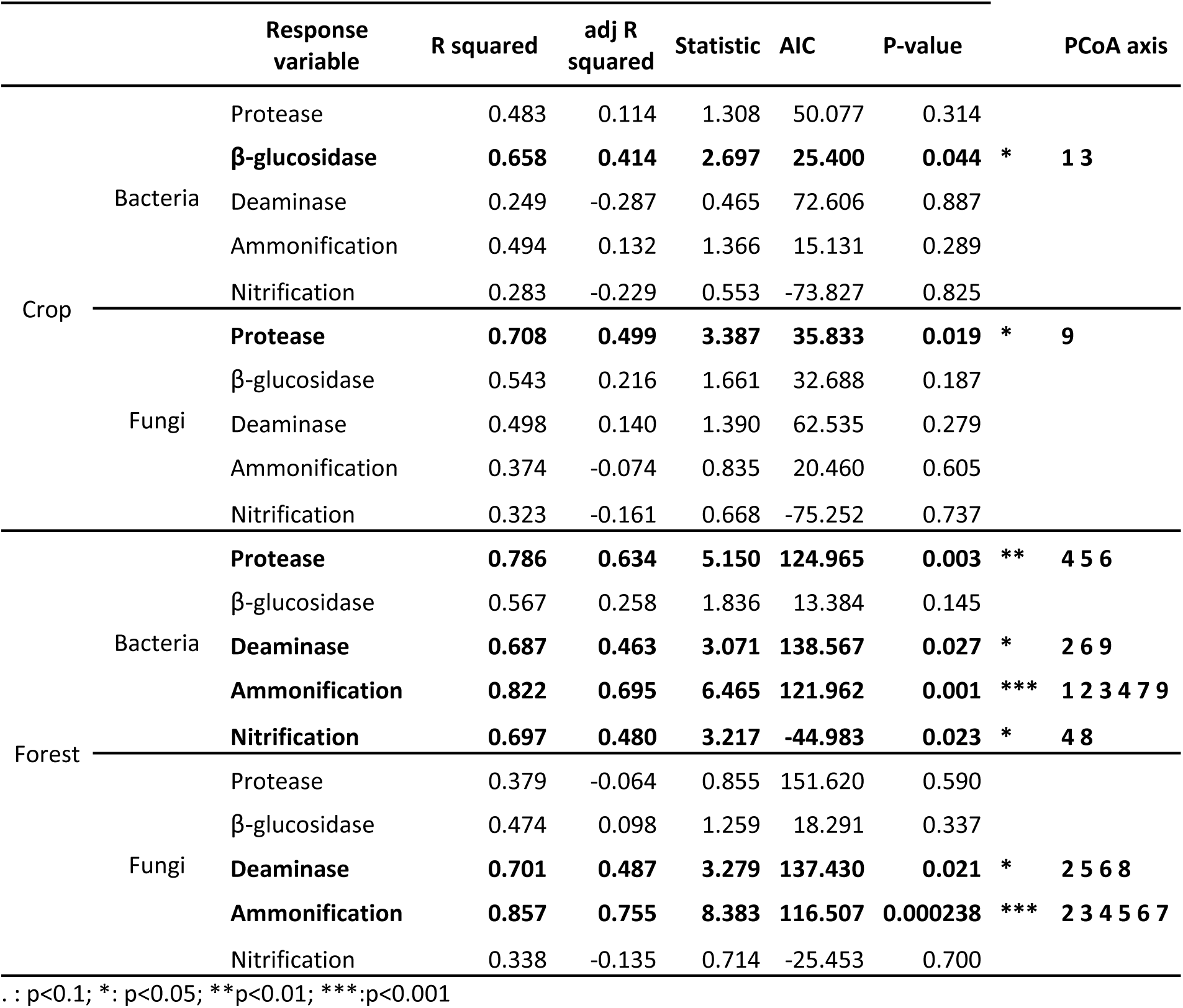
Linear models (LM) results of enzymatic activities (protease, β-glucosidase and deaminase), ammonification and nitrification associated with bacterial and fungal communities’ structure (PcoA axes) in the synthetic soil.

For the depolymerization capacities and mineralization processes that were significantly explained by microbial community structure and composition, we looked for positive

Spearman correlations with individual ASVs (Table S3 Fig S.1; S.2). For bacteria, we found multiple ASVs belonging to the *Paenibacillus* genus. Specifically, ASV # 228 in crop soil correlated with β-glucosidase, ASVs #1829 and #1952 in forest soil correlated with deaminase and ASVs #397 and #360 correlated with ammonification in the forest soil. For fungi, we found ASVs belonging to *Eurotiomycetes* and *Leotiomycetes* correlated to deaminase activity and ASVs belonging to *Tremellomycetes*, *Sordariomycetes* and *Dothideomycetes* correlated with ammonification. Many of the correlations found highlighted ASVs that were absent in most samples, and relatively highly abundant in a few samples that also had high process rates, suggesting that the presence of these ASVs was potentially important for these processes.

## 4. Discussion

We wanted to understand the link between soil microbial alpha diversity and microbial necromass depolymerization and mineralization. We successfully created stable microbial communities with distinct levels of alpha diversity by diluting crop and forest soil microbial communities. However, in contrast to our hypothesis, we did not find strong links between alpha diversity and potential depolymerization enzymatic activities or mineralization (ammonification and nitrification). The only significant effects we found were higher enzymatic activities at some intermediate dilutions, without a clear trend.

The soil processes varied substantially between replicates of the same alpha diversity level, especially in the lower diversity levels. This variation was mirrored in beta dispersion, with microbial communities of the less diversified samples varying more than the more diversified samples. This prompted us to explore if the variation in beta diversity could explain the variation in processes, and we did find that beta diversity, in general, and more specifically the abundance of some ASVs were linked to depolymerization and mineralization processes, especially from the forest communities in the synthetic soil. Overall, we found that beta diversity, and not alpha diversity, was more likely to explain depolymerization and subsequent ammonification and nitrification from crop and forest microbial communities. This is well aligned with our previous study where shifts in microbial community composition induced by differences in crop rotation resulted in different ammonium content in the rhizosphere of the following crop (L’Espérance et al., 2025). Accordingly, the composition of bacterial communities (i.e., beta diversity) was previously shown to impact β-glucosidase activity more than taxonomic diversity (i.e., alpha diversity) (Bailey et al., 2013).

We were able to maintain distinct levels of bacterial and fungal diversity throughout the two phases of our experiment. The dilution to extinction method creates communities with a gradient of diversity (Dıaz et al., 2003; Philippot et al., 2013) through the loss of rare species as dilution increases (Hol et al., 2010). This technique has been shown to impact narrow processes like nitrification and denitrification (Philippot et al., 2013; King et al., 2023). In our case, probably because depolymerization is a broad process mediated by different exoenzymes from diverse microbes, we found few significant differences between depolymerization capacity of the different dilutions (i.e., different alpha diversity levels). When we did find an effect of alpha diversity on the process measured, it was related to increased enzymatic activities (β-glucosidase in native forest soil, in synthetic crop soil and protease in synthetic forest soil) or decreased mineralization (nitrification in synthetic crop soil) at intermediate dilutions (i.e., intermediate diversity levels). These observations were probably related to a lowered competition at these diversity levels. Indeed, higher diversity can create competition for resources (Langenheder et al., 2010). As C and N depolymerization is based on the production of extracellular enzymes, this process could also be viewed through the lens of “social cheating” (Smith and Schuster, 2019). Since exoenzymes are not bound to their producers, bioavailable C and N resulting from depolymerization can be used by other non-producing microorganisms (Smith and Schuster, 2019). Allison and Folse (2012) showed with a model that at high diversity, depolymerization declines due to lower levels of enzyme production (Folse III and Allison, 2012). Higher microbial diversity is associated with higher competition, and, on the contrary, a lower microbial diversity leads to the loss of key species in exoenzymes production. Therefore, we had expected that, like the ‘goldilocks’ principle states, the intermediate diversity would have been the best, since there are plenty of exoenzyme producers and less competition and fewer cheaters (Berlow et al., 2008). But the goldilocks principle was only found to be true for the C associated exoenzyme (β-glucosidases), whereas N associated exoenzymes (proteases and deaminases) did not respond to shifts in alpha diversity.

Another explanation for the higher process rates that were measured at intermediate diversity is the stochastic variation in the presence of proficient microorganisms induced by diluting the soils. Since the dilution to extinction technique is based on the random removal of microbes (Yan et al., 2015), this can results in higher stochasticity in community composition, especially at higher dilutions (Riddley et al., 2025), as we observed in our data (both in native and synthetic soil). Our correlation analysis showed that some ASVs in the synthetic soil were absent from most soils and then relatively abundant in a few soils, which also had high process rates. Among those, protease potential activity (forest soil) was explained by bacterial community structure and two *Rhizobium* ASVs – a genus known to produce extracellular proteases (Gopalakrishnan et al., 2015) – were positively correlated with higher protease activity. Deaminase potential activity from forest soil was also explained by bacterial and fungal beta-diversity, suggesting that the two N depolymerization steps measured are linked to microbial community structure and composition. The low diversity communities are also more vulnerable to the influence of the microorganisms that potentially remained after soil sterilization, or that would come from the environment (Van Elsas et al., 2012). Nevertheless, the stochastic establishment of microbial communities in the synthetic and native soil at lower diversity may explain the high variability in N associated exoenzymes potential activities across replicates and the lack of link between N depolymerization and alpha diversity. This suggests, in contrast to our hypothesis, that N depolymerization capacity depends more on microbial community structure and composition than alpha diversity.

In the synthetic soil, different amounts of microbial necromass were added to forest and crop soil microbial communities to match the amount in real conditions (Wang et al., 2021). Higher exoenzyme activities from forest microbial communities are therefore in line with other findings claiming that the production of many exoenzymes is inducible (Geisseler and Horwath, 2008). However, the relatively low and variable enzymatic activity that we observed could be linked to the relatively low amount of rapidly usable C and N for enzyme production (Allison and Vitousek, 2005). We could see interesting differences in mineralization rates between the two soils. At intermediate dilutions, there was less necromass mineralization from crop soil microbial communities and more necromass mineralization from forest soil microbial communities. A possible explanation for this opposite effect of dilution on the mineralization by forest and crop communities could be linked to their adaptation to their environment.

Crop soils are usually fertilized and enriched in easily available nutrients, selecting for copiotrophs that do not produce exoenzymes to scavenge nutrients (Wang et al., 2024). In contrast, forest soils are more often nutrient-limited, and nutrients are mostly contained in the soil organic matter, which selects for microbes producing exoenzymes (Sinsabaugh and Follstad Shah, 2012; Waring et al., 2014). Crop soil mineralization rate probably decreased at intermediate microbial diversity because there are fewer microbes capable to use necromass-derived N. In lower dilution, mineralization rate is higher probably due to the stochastic selection of microorganisms able to use necromass-derived N and to mineralize. In contrast, mineralization rate in lower dilution increased in forest soil due to a reduced competition.

Another plausible explanation is the lower fungal:bacterial ratio of crop soil than forest soil (Djemiel et al., 2023). Fungi have a lower N demand than bacteria, with a C:N ratio between 5:1 and 15:1 for fungi compared to 3:1 and 6:1 for bacteria (Strickland and Rousk, 2010). Fungi thus need less N for their growth and communities containing more fungi than bacteria could release more N when mineralizing microbial necromass (Högberg et al., 2007). In line with this, we found a *Tremellomycetes* (ASV 8), two *Sordariomycetes* (ASV 80 and 810) and a *Dothiodeomycetes* (ASV 341), positively correlated with higher ammonification rate from forest soil.

Fungi and bacteria did not respond to the dilution similarly. This could be related to their size as Farjalla et al. (2012) showed that the role of stochastic processes in microbial community assembly decreases with size (Farjalla et al., 2012). Fungi also form multicellular hyphae, which are not affected in the same way as bacterial cells during the dilution process (Collado et al., 2007). In addition, the difference in the nature of the amplicon used could also have an impact on our observations. The ITS regions are non-coding DNA sequences in the fungal genome and are typically more variable than coding ribosomal regions (Schoch et al., 2012), resulting in a different taxonomic resolution of the fungal vs. bacterial ASVs. Intriguingly, fungal alpha diversity in the synthetic soil seemed to mirror β-glucosidase potential activity; both measures reached their maximum in D4 for the crop soil and at D2 for the forest soil. The enzyme β-glucosidase is crucial to depolymerize large carbon compounds coming from cellulose (Alef and Nannipieri, 1995). More specifically, it cleaves the β-1,4 bond of β-glucoside (Cañizares et al., 2011). In the synthetic soil, there was no cellulose nor plant necromass, but there was β-glucosidase potential activity measured. This suggests that microbes produce β-glucosidase without the presence of plant necromass. Some β-glucosidase can degrade other sugar polymers (Ketudat Cairns and Esen, 2010; Finley et al., 2025).

## 5. Conclusion

In this work, we showed that C and N associated exoenzymes potential activities is generally not affected by microbial alpha diversity. The soil dilution approach we used led to an increase variability in community composition with decreasing alpha diversity. Some of these communities with a similar alpha diversity were proficient exoenzymes producers and microbial necromass mineralizers, whereas other were not, which explains why alpha diversity *per se* was not a significant factor. Rather, beta diversity and the abundance of specific microorganisms explained more significantly the patterns observed in soil processes, especially for the forest soil community. Thus, efforts to increase soil nitrogen supply for crops should not necessarily target an indiscriminate increase in soil alpha diversity but focus on activating specific microbial taxa. Specifically, we think that targeting specific, proficient exoenzyme producers could lead to more N release trough depolymerization and mineralization for crops. More generally, our results also suggest that positive relationships between alpha diversity and ecosystem processes might not always hold true and could be influence by community composition.

## 6. Funding Sources

This work was supported by the FRQNT Programme de recherche en partenariat (grant number 322604), the Fondation de l’université du Québec en Abitibi-Témiscamingue and the Entente sectorielle bioalimentaire de l’Abitibi-Témiscamingue. ELE was supported by a Fondation Armand-Frappier doctoral scholarship.

## Acknowledgements

We extend our gratitude to our interns: Raphaël Boisvert and Hubert Côté, whose hard work in the lab, particularly with DNA extractions, significantly contributed to this project. We are thankful to Johanne Massy for sampling and sending the soil used in our experiment. We also thank our colleagues for their help, support, and ideas throughout this project.

---

**Table S1:** PERMANOVA results of Hellinger-transformed Euclidian distances from bacterial and fungal communities (16S and ITS) data in their original soil.

| Microbes | Soil | Variable | Df | SumOfSqs | R2 | F | Pr(>F) |
| --- | --- | --- | --- | --- | --- | --- | --- |
| Bacteria | Crop | **Dilution** | **5** | **7.595** | **0.350** | **2.585** | **0.001 \*\*** |
|  |  | Residual | 24 | 14.101 | 0.650 |  |  |
|  |  | Total | 29 | 21.696 | 1.000 |  |  |
|  | Forest | **Dilution** | **5** | **7.804** | **0.353** | **2.614** | **0.001 \*\*** |
|  |  | Residual | 24 | 14.333 | 0.647 |  |  |
|  |  | Total | 29 | 22.137 | 1.000 |  |  |
| Fungi | Crop | **Dilution** | **5** | **6.618** | **0.298** | **2.042** | **0.001 \*\*** |
|  |  | Residual | 24 | 15.559 | 0.702 |  |  |
|  |  | Total | 29 | 22.176 | 1.000 |  |  |
|  | Forest | **Dilution** | **5** | **9.643** | **0.484** | **4.503** | **0.001 \*\*** |
|  |  | Residual | 24 | 10.278 | 0.516 |  |  |
|  |  | Total | 29 | 19.921 | 1.000 |  |  |

\*: p<0.05; \*\*:p<0.01; \*\*\*:p<0.001
Df, SumOfSqs, R2, F and p-value in bold are the significant values (p<0.05)

**Table S2:** PERMANOVA results of Hellinger-transformed Euclidian distances from bacterial and fungal communities (16S and ITS) data in the synthetic soil.

| Microbes | Soil | Variable | Df | SumOfSqs | R2 | F | Pr(>F) |  |
| --- | --- | --- | --- | --- | --- | --- | --- | --- |
| Bacteria | Crop | **Dilution** | **5** | **8.441** | **0.435** | **3.698** | **0.001** | **\*\*** |
|  |  | Residual | 24 | 10.957 | 0.565 |  |  |  |
|  |  | Total | 29 | 19.398 | 1 |  |  |  |
|  | Forest | **Dilution** | **5** | **7.681** | **0.350** | **2.588** | **0.001** | **\*\*** |
|  |  | Residual | 24 | 14.243 | 0.650 |  |  |  |
|  |  | Total | 29 | 21.924 | 1 |  |  |  |
| Fungi | Crop | **Dilution** | **5** | **7.187** | **0.327** | **2.330** | **0.001** | **\*\*** |
|  |  | Residual | 24 | 14.803 | 0.673 |  |  |  |
|  |  | Total | 29 | 21.990 | 1 |  |  |  |
|  | Forest | **Dilution** | **5** | **10.531** | **0.466** | **4.193** | **0.001** | **\*\*** |
|  |  | Residual | 24 | 12.056 | 0.534 |  |  |  |
|  |  | Total | 29 | 22.587 | 1 |  |  |  |

\*: p<0.05; \*\*:p<0.01; \*\*\*:p<0.001
Df, SumOfSqs, R2, F and p-value in bold are the significant values (p<0.05)

**Table S3:**
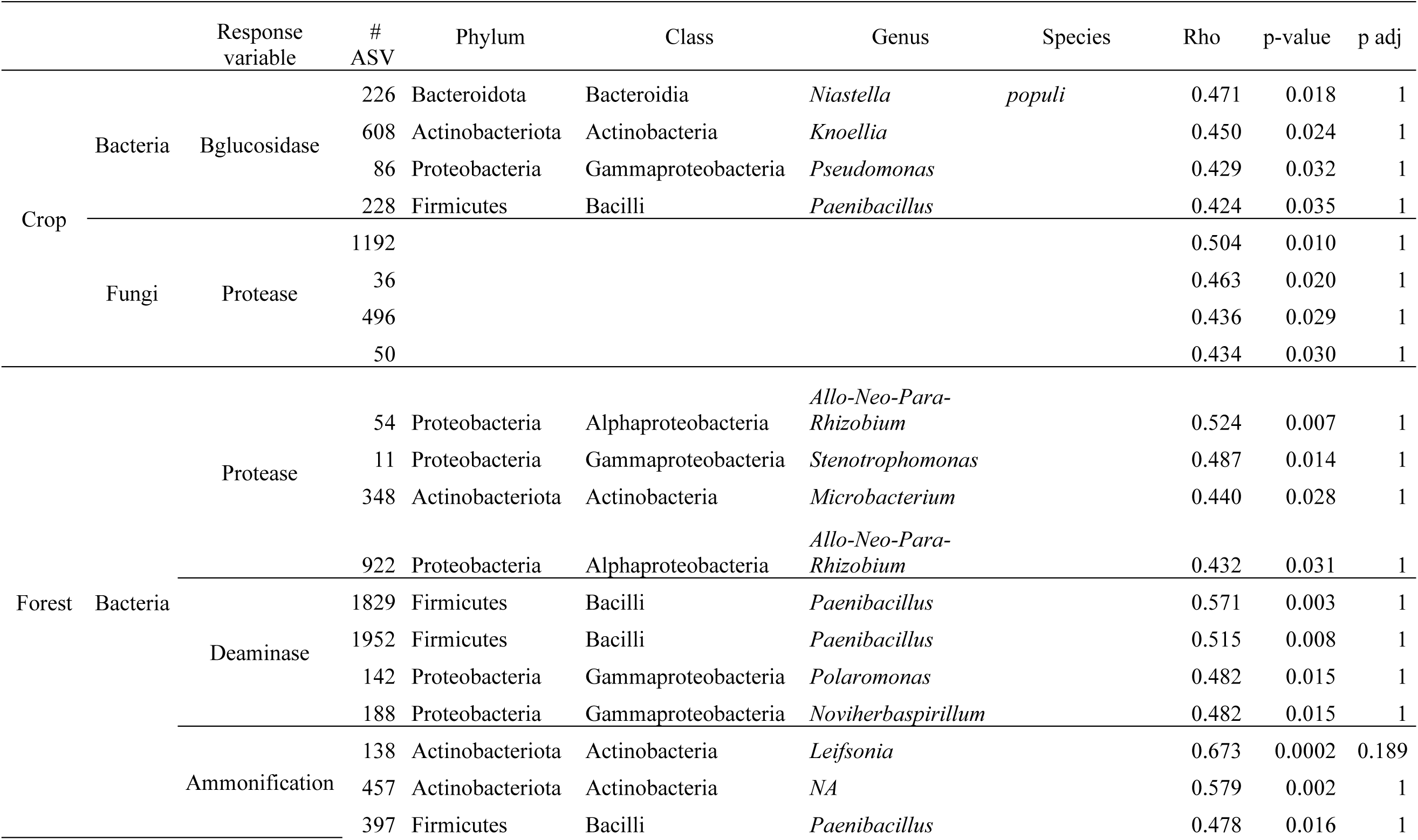
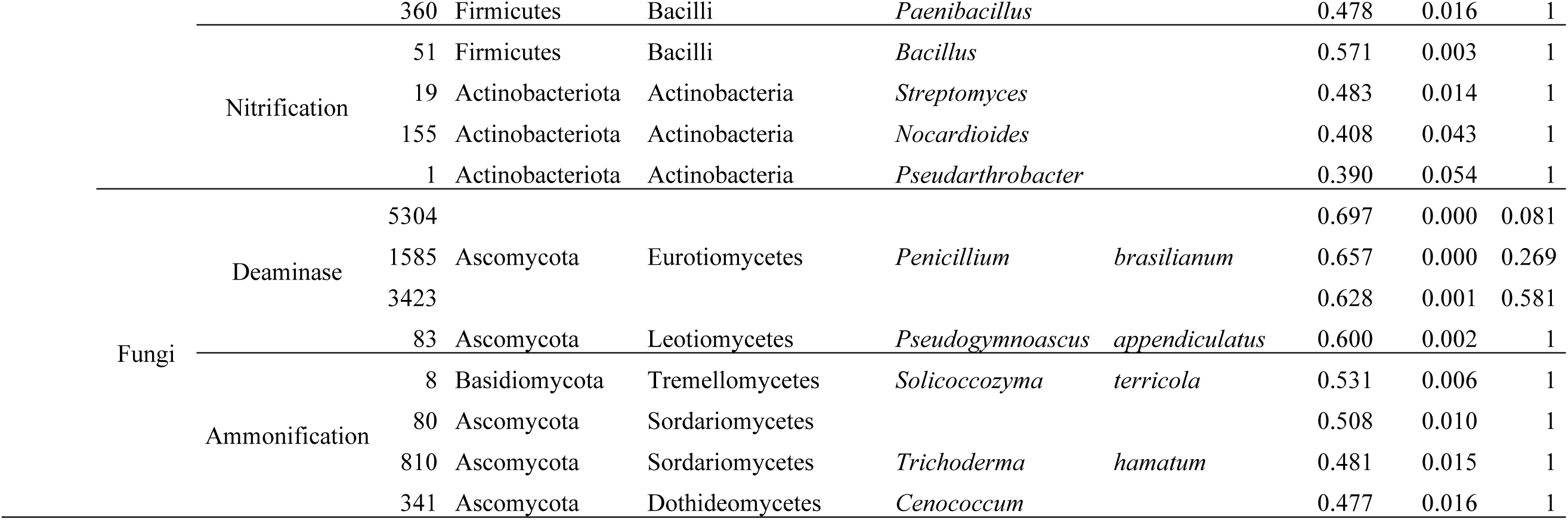
ASVs positively correlated with depolymerization capacities and mineralization processes that was significantly explained by microbial communities’ structure and composition.

**Figure S1:**
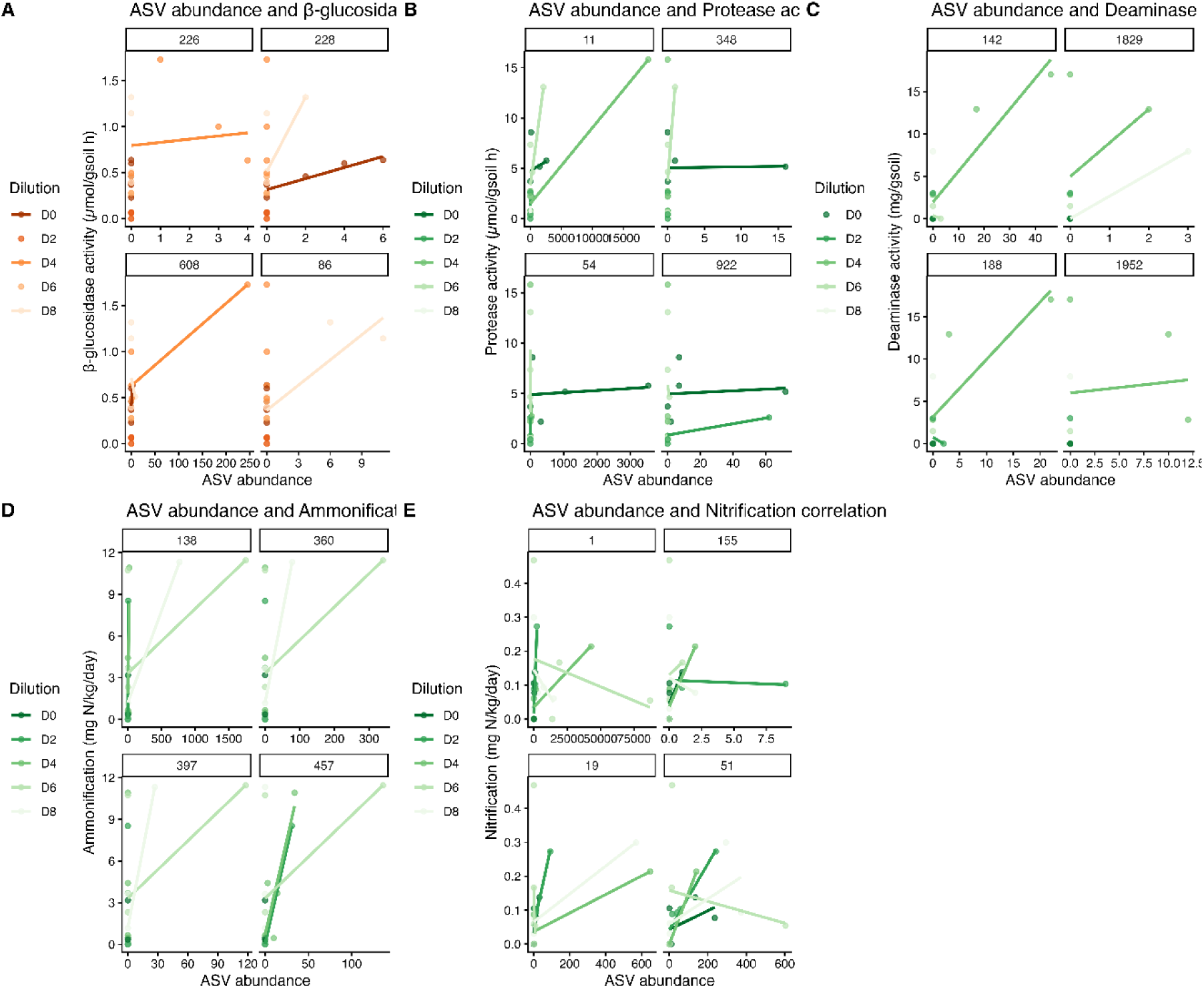
Bacterial ASVs positively correlated to exoenzymes potential activities (protease, β-glucosidase and deaminase) and mineralization processes explained by beta-diversity (PCoA axis) in the synthetic soil. ASVs found in crop communities are in red and ASVs found in forest communities are in green.

**Figure S2:**
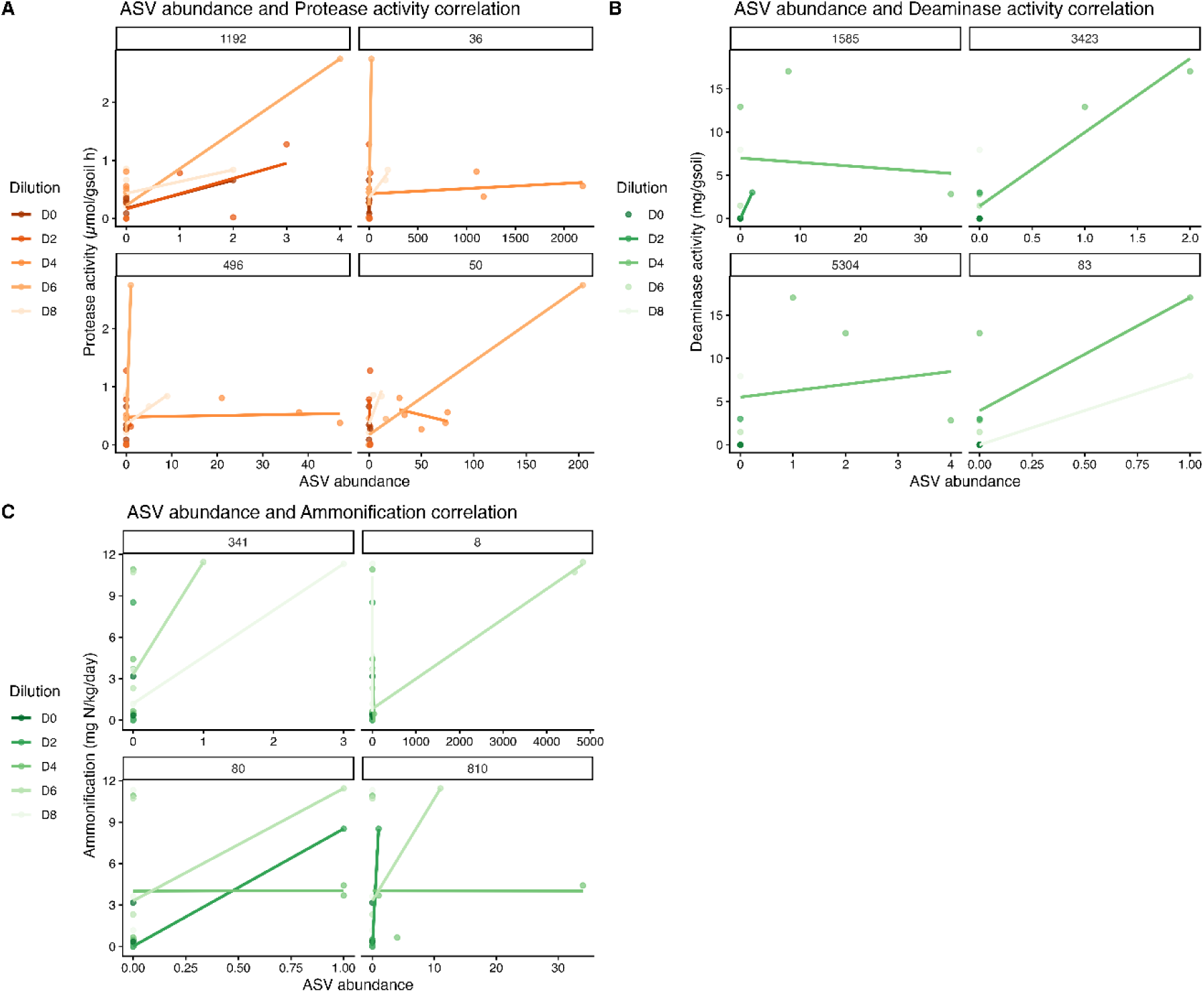
Fungal ASVs positively correlated to exoenzymes potential activities (protease, β-glucosidase and deaminase) and mineralization processes explained by beta-diversity (PCoA axis) in the synthetic soil. ASVs found in crop communities are in red and ASVs found in forest communities are in green.

## Notes

### Competing Interest Statement

The authors have declared no competing interest.

## References

Alef, K., Nannipieri, P., 1995. Methods in applied soil microbiology and biochemistry.

Allison, S.D., Vitousek, P.M., 2005. Responses of extracellular enzymes to simple and complex nutrient inputs. Soil Biology and Biochemistry 37, 937–944.

Bailey, V.L., Fansler, S.J., Stegen, J.C., McCue, L.A., 2013. Linking microbial community structure to β-glucosidic function in soil aggregates. The ISME journal 7, 2044–2053.

Baumann, K., Dignac, M.-F., Rumpel, C., Bardoux, G., Sarr, A., Steffens, M., Maron, P.-A., 2013. Soil microbial diversity affects soil organic matter decomposition in a silty grassland soil. Biogeochemistry 114, 201–212.

Berlow, E.L., Brose, U., Martinez, N.D., 2008. The “Goldilocks factor” in food webs. Proceedings of the National Academy of Sciences 105, 4079–4080.

Buckeridge, K.M., Mason, K.E., Ostle, N., McNamara, N.P., Grant, H.K., Whitaker, J., 2022. Microbial necromass carbon and nitrogen persistence are decoupled in agricultural grassland soils. Communications Earth & Environment 3, 114.

Callahan, B.J., McMurdie, P.J., Rosen, M.J., Han, A.W., Johnson, A.J.A., Holmes, S.P., 2016. DADA2: High-resolution sample inference from Illumina amplicon data. Nature methods 13, 581–583.

Cañizares, R., Benitez, E., Ogunseitan, O.A., 2011. Molecular analyses of β-glucosidase diversity and function in soil. European Journal of Soil Biology 47, 1–8.

Chen, H., Ma, K., Lu, C., Fu, Q., Qiu, Y., Zhao, J., Huang, Y., Yang, Y., Schadt, C.W., Chen, H., 2022. Functional redundancy in soil microbial community based on metagenomics across the globe. Frontiers in microbiology 13, 878978.

Collado, J., Platas, G., Paulus, B., Bills, G.F., 2007. High-throughput culturing of fungi from plant litter by a dilution-to-extinction technique. FEMS Microbiology Ecology 60, 521–533.

Dıaz, S., Symstad, A.J., Chapin, F.S., Wardle, D.A., Huenneke, L.F., 2003. Functional diversity revealed by removal experiments. Trends in ecology & evolution 18, 140–146.

Djemiel, C., Dequiedt, S., Bailly, A., Tripied, J., Lelièvre, M., Horrigue, W., Jolivet, C., Bispo, A., Saby, N., Valé, M., 2023. Biogeographical patterns of the soil fungal: bacterial ratio across France. Msphere 8, e00365–00323.

Farjalla, V.F., Srivastava, D.S., Marino, N.A., Azevedo, F.D., Dib, V., Lopes, P.M., Rosado, A.S., Bozelli, R.L., Esteves, F.A., 2012. Ecological determinism increases with organism size. Ecology 93, 1752–1759.

Finley, B.K., Enalls, B.C., de Raad, M., Al Said, M., Chen, M., Joyner, D.C., Hazen, T.C., Northen, T.R., Chakraborty, R., 2025. Unraveling the influence of microbial necromass on subsurface microbiomes: metabolite utilization and community dynamics. ISME communications 5, ycaf006.

Folse III, H.J., Allison, S.D., 2012. Cooperation, competition, and coalitions in enzyme-producing microbes: social evolution and nutrient depolymerization rates. Frontiers in microbiology 3, 338.

Geisseler, D., Horwath, W.R., 2008. Regulation of extracellular protease activity in soil in response to different sources and concentrations of nitrogen and carbon. Soil Biology and Biochemistry 40, 3040–3048.

Giller, K., Beare, M., Lavelle, P., Izac, A.-M., Swift, M., 1997. Agricultural intensification, soil biodiversity and agroecosystem function. Applied Soil Ecology 6, 3–16.

Gopalakrishnan, S., Sathya, A., Vijayabharathi, R., Varshney, R.K., Gowda, C.L., Krishnamurthy, L., 2015. Plant growth promoting rhizobia: challenges and opportunities. 3 Biotech 5, 355–377.

Guenet, B., Leloup, J., Hartmann, C., Barot, S., Abbadie, L., 2011. A new protocol for an artificial soil to analyse soil microbiological processes. Applied Soil Ecology 48, 243–246.

Heemsbergen, D., Berg, M., Loreau, M., Van Hal, J., Faber, J., Verhoef, H., 2004. Biodiversity effects on soil processes explained by interspecific functional dissimilarity. Science 306, 1019–1020.

Högberg, M.N., Chen, Y., Högberg, P., 2007. Gross nitrogen mineralisation and fungi-to-bacteria ratios are negatively correlated in boreal forests. Biology and Fertility of Soils 44, 363–366.

Hol, W.G., De Boer, W., Termorshuizen, A.J., Meyer, K.M., Schneider, J.H., Van Dam, N.M., Van Veen, J.A., Van Der Putten, W.H., 2010. Reduction of rare soil microbes modifies plant–herbivore interactions. Ecology letters 13, 292–301.

Jan, M.T., Roberts, P., Tonheim, S.K., Jones, D.L., 2009. Protein breakdown represents a major bottleneck in nitrogen cycling in grassland soils. Soil Biology and Biochemistry 41, 2272–2282.

Ketudat Cairns, J.R., Esen, A., 2010. β-Glucosidases. Cellular and Molecular Life Sciences 67, 3389–3405.

King, W.L., Richards, S.C., Kaminsky, L.M., Bradley, B.A., Kaye, J.P., Bell, T.H., 2023. Leveraging microbiome rediversification for the ecological rescue of soil function. Environmental Microbiome 18, 1–13.

L’Espérance, E., Poirier, V., Lavergne, S., Yergeau, É., 2025. Previous legume identity influences wheat protein content through organic matter depolymerization. bioRxiv, 2025.2012. 2017.694934.

Langenheder, S., Bulling, M.T., Solan, M., Prosser, J.I., 2010. Bacterial biodiversity-ecosystem functioning relations are modified by environmental complexity. PLoS One 5, e10834.

Loreau, M., 2004. Does functional redundancy exist? Oikos 104, 606–611.

Nguyen, T.T., Myrold, D.D., Mueller, R.S., 2019. Distributions of extracellular peptidases across prokaryotic genomes reflect phylogeny and habitat. Frontiers in microbiology 10, 413.

Philippot, L., Spor, A., Hénault, C., Bru, D., Bizouard, F., Jones, C.M., Sarr, A., Maron, P.-A., 2013. Loss in microbial diversity affects nitrogen cycling in soil. The ISME journal 7, 1609–1619.

Pold, G., Saghaï, A., Jones, C.M., Hallin, S., 2025. Denitrification is a community trait with partial pathways dominating across microbial genomes and biomes. Nature Communications 16, 9495.

Riddley, M., Hepp, S., Hardeep, F., Nayak, A., Liu, M., Xing, X., Zhang, H., Liao, J., 2025. Differential roles of deterministic and stochastic processes in structuring soil bacterial ecotypes across terrestrial ecosystems. Nature Communications 16, 2337.

Schimel, J.P., Bennett, J., 2004. Nitrogen mineralization: challenges of a changing paradigm. Ecology 85, 591–602.

Schoch, C.L., Seifert, K.A., Huhndorf, S., Robert, V., Spouge, J.L., Levesque, C.A., Chen, W., Consortium, F.B., List, F.B.C.A., Bolchacova, E., 2012. Nuclear ribosomal internal transcribed spacer (ITS) region as a universal DNA barcode marker for Fungi. Proceedings of the National Academy of Sciences 109, 6241–6246.

Schulten, H.-R., Schnitzer, M., 1997. The chemistry of soil organic nitrogen: a review. Biology and Fertility of Soils 26, 1–15.

Shao, Y.-H., Lu, H.-P., Wu, J.-H., 2025. Microbial diversity supports nitrification: insights from a full-scale anoxic/oxic wastewater treatment process. Applied and environmental microbiology, e01803–01825.

Simpson, A.J., Simpson, M.J., Smith, E., Kelleher, B.P., 2007. Microbially derived inputs to soil organic matter: are current estimates too low? Environmental Science & Technology 41, 8070–8076.

Sinsabaugh, R., 1994. Enzymic analysis of microbial pattern and process. Biology and Fertility of Soils 17, 69–74.

Sinsabaugh, R.L., Follstad Shah, J.J., 2012. Ecoenzymatic stoichiometry and ecological theory. Annual review of ecology, evolution, and systematics 43, 313–343.

Smith, P., Schuster, M., 2019. Public goods and cheating in microbes. Current biology 29, R442–R447.

Stein, L.Y., Klotz, M.G., 2016. The nitrogen cycle. Curr Biol 26, R94–98.

Strickland, M.S., Rousk, J., 2010. Considering fungal:bacterial dominance in soils – Methods, controls, and ecosystem implications. Soil Biology and Biochemistry 42, 1385–1395.

Szoboszlay, M., Dohrmann, A.B., Poeplau, C., Don, A., Tebbe, C.C., 2017. Impact of land-use change and soil organic carbon quality on microbial diversity in soils across Europe. FEMS Microbiology Ecology 93, fix146.

Van Elsas, J.D., Chiurazzi, M., Mallon, C.A., Elhottovā, D., Krištůfek, V., Salles, J.F., 2012. Microbial diversity determines the invasion of soil by a bacterial pathogen. Proceedings of the National Academy of Sciences 109, 1159–1164.

Waldrop, M., Balser, T., Firestone, M., 2000. Linking microbial community composition to function in a tropical soil. Soil Biology and Biochemistry 32, 1837–1846.

Wanek, W., Mooshammer, M., Blöchl, A., Hanreich, A., Richter, A., 2010. Determination of gross rates of amino acid production and immobilization in decomposing leaf litter by a novel 15N isotope pool dilution technique. Soil Biology and Biochemistry 42, 1293–1302.

Wang, B., An, S., Liang, C., Liu, Y., Kuzyakov, Y., 2021. Microbial necromass as the source of soil organic carbon in global ecosystems. Soil Biology and Biochemistry 162, 108422.

Wang, C., Wang, N., Zhu, J., Liu, Y., Xu, X., Niu, S., Yu, G., Han, X., He, N., 2018. Soil gross N ammonification and nitrification from tropical to temperate forests in eastern China. Functional Ecology 32, 83–94.

Wang, X., Lin, J., Peng, X., Zhao, Y., Yu, H., Zhao, K., Barberán, A., Kuzyakov, Y., Dai, Z., 2024. Microbial rrn copy number is associated with soil C: N ratio and pH under long-term fertilization. Science of the Total Environment 954, 176675.

Waring, B.G., Weintraub, S.R., Sinsabaugh, R.L., 2014. Ecoenzymatic stoichiometry of microbial nutrient acquisition in tropical soils. Biogeochemistry 117, 101–113.

Yan, Y., Kuramae, E.E., Klinkhamer, P.G., van Veen, J.A., 2015. Revisiting the dilution procedure used to manipulate microbial biodiversity in terrestrial systems. Applied and environmental microbiology 81, 4246–4252.

Yoon, J.-H., Adhikari, M., Jeong, S.S., Lee, S.P., Kim, H.S., Lee, G.S., Park, D.H., Kim, H., Yang, J.E., 2024. Microbial diversity of soils under different land use and chemical conditions. Applied Biological Chemistry 67, 111.

Young, I., Ritz, K., 2000. Tillage, habitat space and function of soil microbes. Soil and Tillage Research 53, 201–213.

